# Inferring neural dynamics of memory during naturalistic social communication

**DOI:** 10.1101/2024.01.26.577404

**Authors:** Rich Pang, Christa Baker, Mala Murthy, Jonathan Pillow

## Abstract

Memory processes in complex behaviors like social communication require forming representations of the past that grow with time. The neural mechanisms that support such continually growing memory remain unknown. We address this gap in the context of fly courtship, a natural social behavior involving the production and perception of long, complex song sequences. To study female memory for male song history in unrestrained courtship, we present ‘Natural Continuation’ (NC)—a general, simulation-based model comparison procedure to evaluate candidate neural codes for complex stimuli using naturalistic behavioral data. Applying NC to fly courtship revealed strong evidence for an adaptive population mechanism for how female auditory neural dynamics could convert long song histories into a rich mnemonic format. Song temporal patterning is continually transformed by heterogeneous nonlinear adaptation dynamics, then integrated into persistent activity, enabling common neural mechanisms to retain continuously unfolding information over long periods and yielding state-of-the-art predictions of female courtship behavior. At a population level this coding model produces multi-dimensional advection-diffusion-like responses that separate songs over a continuum of timescales and can be linearly transformed into flexible output signals, illustrating its potential to create a generic, scalable mnemonic format for extended input signals poised to drive complex behavioral responses. This work thus shows how naturalistic behavior can directly inform neural population coding models, revealing here a novel process for memory formation.

---

Many fundamental behaviors depend on the ability to form and update memories of a continually growing past. Responding appropriately in conversation, for instance, may require remembering the history of the conversation over multiple timescales, while efficiently updating one’s internal representation of this history as it unfolds. Such memory-dependent tasks are faced by a variety of flexible intelligent systems (1, 2) and mediate important phenomena like information transmission through social networks (3)—yet how the brain solves this is unknown. While a number of theories can explain a variety of controlled memory experiments (4–8), they do not generalize to more natural settings like uncontrolled social communication. As a result, how biological neural algorithms encode natural input histories like those in communication, what neural processes support such encoding, how representations evolve online to continually incorporate new information, how these encoding processes advantage computation, and whether such processes can be described by general principles, are not understood.

Natural social behaviors represent an excellent system for addressing these questions. During fly (*D. melanogaster*) courtship, males sing complex and variable song sequences to females. Songs are composed of two main syllables or “song modes” called *sine* and *pulse* for their acoustic waveform, and are interleaved with silences (Fig 1A-C) (9–11). Fly courtship song shares key features with more complex signals like speech or language. First, like phonemes, song elements rapidly fluctuate over time; second, songs can extend over timescales much larger (minutes) than their constituent elements (∼30 ms); third, sufficiently long sequences of song elements rarely repeat, similar to how sentences rarely repeat in say, a news story. Thus, understanding song processing in the fly brain may shed light on how extended, fluctuating input signals are processed and stored in more complex neural systems as well.

**Fig. 1.**
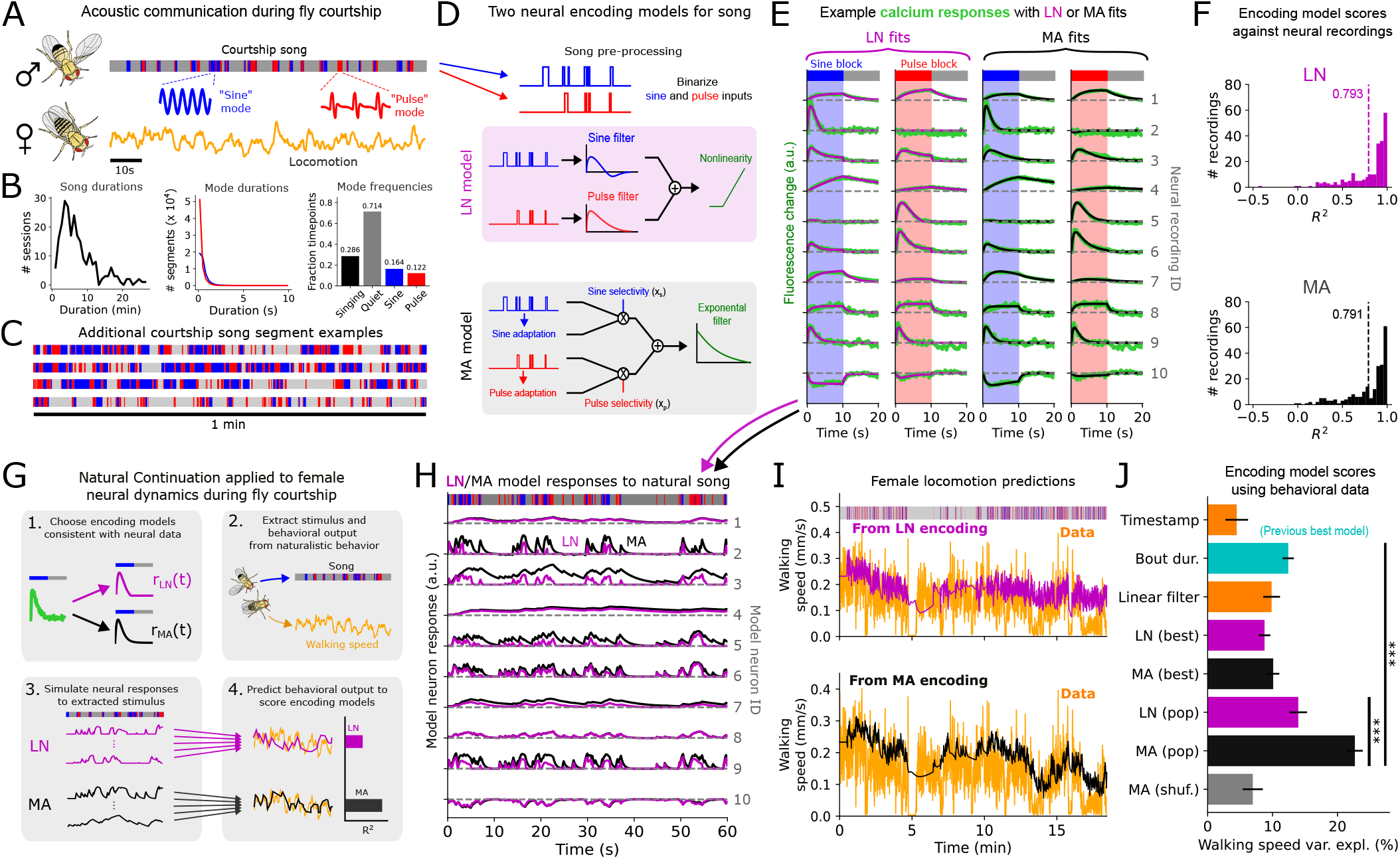
Testing neural encoding models of song history against naturalistic fly courtship data. **A**. Schematic of *Drosophila* acoustic communication, with example song and locomotion trace. Song comprises two main modes termed ‘sine’ and ‘pulse’. **B**. Song feature distributions. **C**. Additional song segment examples from unrestrained courtship. **D**. Schematic of LN and MA encoding models. **E**. Example neural responses measured during calcium imaging experiments (18) in response to a 10s block of either pure sine or pure pulse song, alongside fits with the LN or MA encoding models. Each row shows an example neural recording from one experiment. The calcium responses in the last two columns are a copy of the responses in the first two columns, except overlaid with the MA instead of LN fits. (The last row shows an example neuron whose fluorescence decreased in response to block song.) **F**. *R*^2^ distributions of LN vs MA fits, with mean across recordings indicated by dashed lines. **G**. Schematic of applying Naturalistic Continuation to fly courtship. **H**. Example responses to naturalistic song of either the LN or MA neuron models fit to the block-song responses in E. **I**. Example prediction of female walking speed from a linear readout of the artificial activity produced by either the LN (top) or MA (bottom) model population (224 model neurons total); trace shown is on a held-out courtship session not used to fit the linear readout. **J**. Encoding model scores, computed over held-out sessions across 30 training/test (80/20%) splits of 87 courtship sessions, as well as predictions from the timestamp alone, time-averaged bout duration (averaged over 2 minutes of song history, Fig S14), and a pair of linear filters directly on sine and pulse inputs. Female walking speed was forward-averaged over a 1-second window prior to prediction. “Pop” denotes the entire artificial population was used in fitting the readout; “best” denotes that only the single most behaviorally predictive neuron was used. MA (shuffled) shows the walking speed variance explained from artificial neural activity in response to songs that were shuffled across courtship sessions. Error bars indicate standard error. Stars shown denote P < .0005 (LN (pop) vs MA (pop)) and P < 10^−8^ (MA (pop) vs bout duration) (2-sided t-test).

Here we ask how female neural dynamics encode the history of courtship song to guide memory-dependent locomotor responses. Previous work found that female slowing during courtship could be predicted by the average duration of preceding song bouts (contiguous singing periods), with predictability plateauing at an averaging window of one minute into the past (12). This suggests that the female may remember song features over timescales on the order of minutes, in contrast with the much faster timescales (seconds or less) typically studied in flies (11, 13–16). More recently, it was additionally found that female auditory neural responses to simplified song stimuli are diverse and widespread across her brain (17, 18), suggesting that the memory of song history may be mediated by a rich, multi-dimensional population code. Yet how female auditory neural responses collectively process and possibly store in memory the diverse temporal patterns of natural song to guide her behavior is unknown.

Addressing this question in naturalistic courtship begets a crucial challenge: recording neural activity in animals interferes with their behavior. In flies, for instance, neural activity is recorded only in head-fixed preparations, which may not reflect brain function in natural settings. Recent efforts have therefore focused on analyzing pure-behavioral data with models governed by neurobiological constraints (12, 19), producing models of neural function directly applicable in natural settings. To synthesize and extend these approaches here we elaborate a general procedure—which we term Natural Continuation (NC)—for systematically assessing neural encoding models according to how well they generalize to account for natural behavior. Although we focus on fly courtship, this method can be widely applied to link neural and naturalistic behavioral data to interrogate neural computation in a variety of systems.

Here we apply NC to neural recordings and naturalistic fly courtship data to discover a novel biologically plausible model for female encoding of song history in memory. Song is first processed by a bank of fast nonlinear adaptation processes then integrated into slowly decaying persistent activity. This produces a multi-dimensional neural representation that yields state-of-the-art predictions of female locomotion during courtship via a novel, neurally informed “basis” of interpretable song patterns that predict slowing. We demonstrate the dynamical and functional advantages of this coding model, including near optimal compression of song history by single neurons, together with advection-diffusion-like population dynamics that separate songs over multiple timescales and can be linearly transformed into flexible outputs. Thus, naturalistic behavior data can strongly inform neural coding models for complex stimuli, which improve behavioral predictions, admit sensible mechanistic interpretations, and suggest generalizable algorithms for flexible computation.

## Results

### Natural Continuation

Natural Continuation is a general protocol for testing neural coding models of complex stimuli against naturalistic behavior data. This makes it a widely applicable method for quantitatively combining existing neural recordings with separate behavioral data to infer neural computations in natural settings. Below we describe NC generally, then apply it to fly courtship.

NC comprises 4 key steps (Fig 1G): (1) Using the existing neural data, select a set of candidate neural encoding models mapping stimulus history to neural activity. (2) Choose a naturalistic behavioral dataset from which the stimulus time-series experienced by the animal can be estimated, as well as a behavioral “output” expected to be modulated by the stimulus. (3) Apply each encoding model to the estimated stimulus history at each timepoint to produce a complete artificial neural population recording (one per encoding model) alongside the behavioral data. (4) Score each neural encoding model by the ability of its corresponding artificial recordings to predict the next-timestep behavioral output through a readout.

Iteratively applying NC to specific model classes produces increasingly refined neural encoding models maximally consistent with the behavioral data. Under fairly generous conditions (see Discussion), this yields a moment-to-moment stimulus-to-behavior model mediated by realistic neural population dynamics, together with artificial neural population recordings that can be studied in their own right to shed light on how neural dynamics processes stimuli in natural settings.

### Two neural encoding models for song history

We applied NC to understand how female flies neurally encode male song history during courtship. To implement step (1) we examined auditory neural recordings (18) from head-fixed female flies in response to restricted “block-song” stimuli, i.e. 10-second blocks of either pure sine or pure pulse song (Fig 1E). Recordings from 50 cell types were performed via two-photon imaging of calcium sensor GCaMP6s (20) in response to the block-song stimuli (randomized and interleaved with quiet blocks). Trial-averaged responses from different neurons exhibited diverse temporal profiles (Fig 1E, S1), varying in their song-mode preference and integration and adaptation timescales. A related pan-neuronal imaging experiment (17) at a lower frame rate (2 Hz in (17) vs 8.5 Hz in (18)) yielded a similar response diversity (Fig S1). The neural data thus suggest specific constraints on the shapes of neural responses to simple song stimuli. However, the block-song stimuli presented in these experiments do not reflect natural song, which contains much shorter song modes interleaved in highly variable temporal patterns (Fig 1A-C).

To gain insight into how female neural dynamics encode natural song we considered two types of models for mapping generic song history to female neural activity (Fig 1D). In the linear-nonlinear (LN) model, the response *r*(*t*) of a single model neuron was given by:

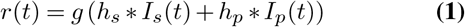

where *I*_*s*_ and *I*_*p*_ are binarized representations of sine and pulse song; *h*_*s*_ and *h*_*p*_ neuron-specific linear filters for sine and pulse song, * represents convolution, and *g* is a signed-rectification rectifying nonlinearity. The LN model reflects a canonical feature-detection computation used to model neural responses in a variety of sensory systems (21–23).

In the “multiplicative adaptation” (MA) model, inspired by models of adaptive neural coding (24–27), a single model neuron’s response *r*(*t*) was given by a simple 4-parameter dynamical system. Specifically:

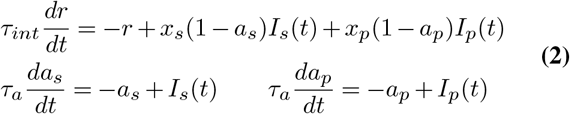

where *τ*_*int*_ and *τ*_*a*_ are integration and adaptation timescales of the neuron and *x*_*s*_ and *x*_*p*_ are its selectivities for sine and pulse song. Intuitively, when either sine or pulse persists for ∼ *τ*_*a*_, its adaptation variable *a*_*s*_ or *a*_*p*_ rises, temporarily diminishing the effect of that song mode input. A notable feature of the MA model useful for memory is that when *τ*_*int*_ is large, this encoding model integrates a nonlinear function of song. At a population level, this geometrically separates song temporal patterns before effectively storing them in persistent activity—this is in contrast to the LN model, in which the nonlinearity is applied only after linear filtering/integration.

We fit each of the two models to each neural recording. To ensure a fair comparison between the two models we derived an analytical formula allowing us to exactly parameterize the LN model with the same 4 parameters as the MA model (See Supplement).

We found that both encoding models were expressive enough to reproduce most of the neural responses to the block-song responses (Fig 1E-F, S2). The LN and MA models differed by less than 1% in the amount of total neural variability they were able to explain across the population. The only neurons poorly fit were those exhibiting “sine-offset” responses (Fig S3), which we discuss later. Thus, the LN and MA models, representing two competing hypotheses for encoding song history, both fit the neural data well and could not be distinguished from the neural data alone.

### The MA encoding best predicts female locomotion

To apply step (2) of NC—in order to compare the LN vs MA encodings of natural song—we used a pure behavioral dataset of naturalistic courtship interactions (11). We used male song as the stimulus and female walking speed as the output. This allowed us to apply step (3) of NC, in which we used the LN and MA encoding models to generate artificial recordings of song-evoked female neural population activity alongside the courtship sessions. Although their block-song responses were nearly indistinguishable, in general the LN and MA responses to naturalistic songs differed, with the LN responses fluctuating rapidly and the MA neurons often exhibiting slower, accumulator-like activity (Fig 1H). Finally, we applied step (4) of NC, scoring the encoding models by how well their neural representations of song history (up till time *t*) could predict female walking speed (time-averaged from *t* till *t*+ 1s) through a linear readout, quantified by walking speed variance explained in held-out courtship sessions.

The LN and MA population encoding models yielded different predictions of female locomotion during courtship (Fig 1I), with the MA model significantly outperforming the LN code (Fig 1J). The MA population also outperformed predictions from time-averaged song bout duration—the best previously conjectured predictor of female walking during courtship (12). While (12) showed that average bout duration could also be computed by single neurons via an adaptive neural mechanism, the collective coding of song by a heterogeneous population had not been probed. Indeed, using only the single best MA neuron yielded similar performance to bout duration, but both were less predictive than the full MA population (Fig 1J). Shuffling songs across courtship sessions strongly degraded performance, indicating that the model-generated activity was predicting locomotion by encoding song history. The same qualitative patterns also held when we predicted female walking speed averaged from *t* to *t* + 1 minute, and when predicting forward or lateral velocity (though these were slightly less predictable than walking speed) (Fig S4), hence our results were largely invariant to the behavioral observables used to score the encoding models. Thus, the naturalistic courtship data is most consistent with an MA population code for song history.

### The MA code is supported by fast adaptation and slow integration

A central feature of robust memory is the ability to keep track of input history over the entire course of a behavior. To assess whether and how the female fly retained song history information over long timescales we iteratively applied NC to perturbed versions of the MA population. For each perturbation we re-simulated neural population activity throughout the courtship sessions, then recomputed female walking speed variance explained by a linear readout (Fig 2A, S4). This first revealed the necessity of pulse selectivity (*x*_*p*_ > 0) and adaptation (*τ*_*a*_ < ∞), as removing either from the population model substantially decreased female walking speed variance explained. The lack of necessity of sine selectivity may reflect redundancy in the population code caused by the high sine-pulse correlation in song (Fig S5).

**Fig. 2.**
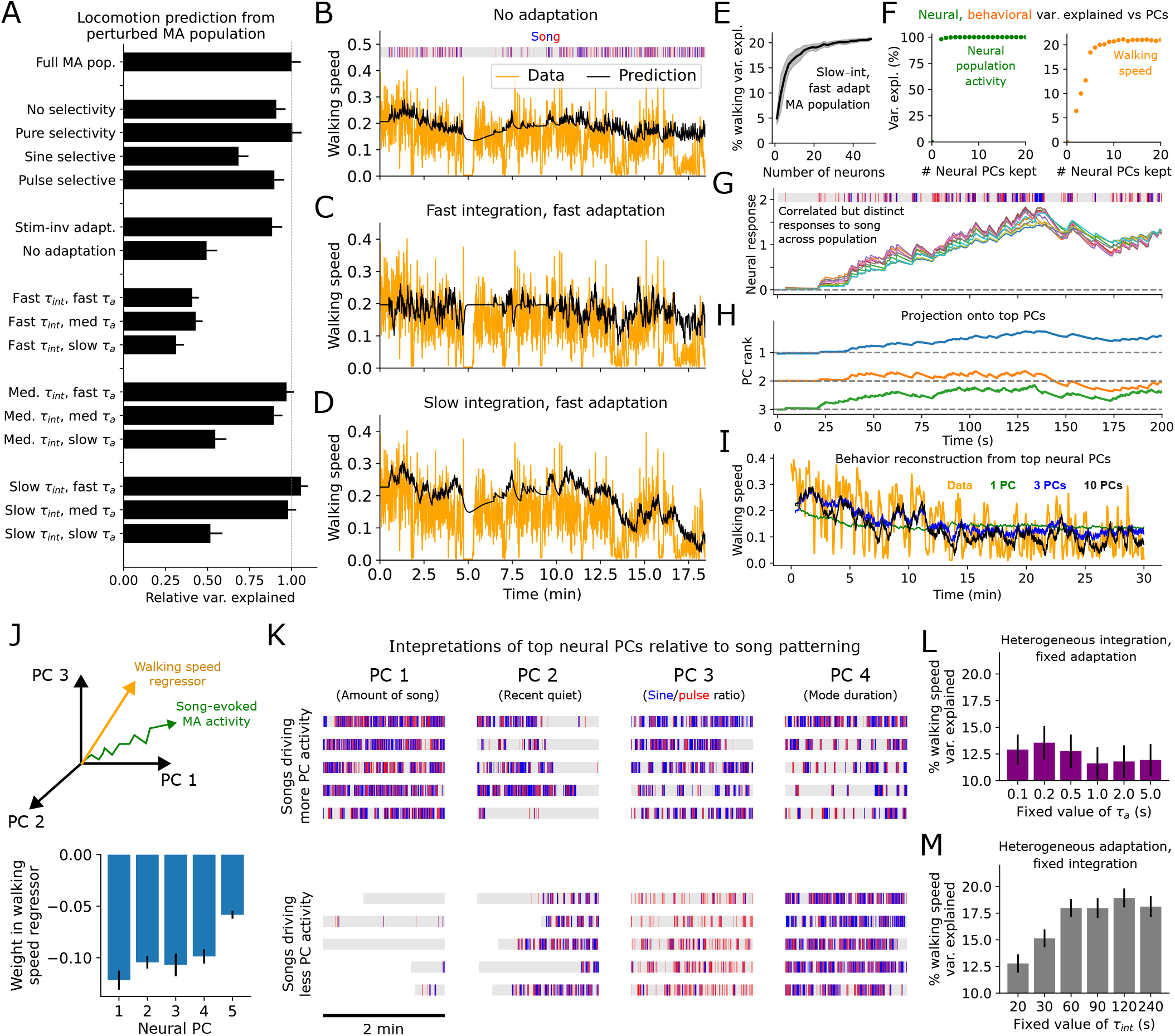
Key features of the MA code for song history. **A**. Predictions of female walking speed from artificial neural recordings generated by perturbed MA population encoding models. Error bars are as in Fig 1J. Here and elsewhere “fast” corresponds to 100*ms* ≤ *τ* < 2*s*, “medium” to 2*s* ≤ *τ* < 20*s*, and “slow” to 20*s* ≤ *τ* < 120*s*. **B**. Female walking speed trace and prediction from an artificial MA population recording with no adaptation. **C**. As in B but from a population with only fast integration and adaptation timescales. **D**. As in B,C except from a population with only slow integration and fast adaptation. **E**. Female walking speed variance explained in held-out sessions from activity produced by a small population of MA neurons with parameters randomly sampled from the fast-adapt/slow-integrate regime, averaged over 30 training/test splits, as a function of the number of neurons included in the population. Shading indicates standard deviation across 30 random instantiations of this MA population. **F**. Left: total neural variance explained (for a population of 20 fast-adapt/slow-integrate MA neurons) vs number of neural PCs kept. Right: female walking speed variance explained vs number of neural PCs kept from the same 20-neuron population. **G**. 10 neural responses out of a 20-neuron population of fast-adapt/slow-integrate MA neurons to example song. **H**. Projection of population activity in G onto top 3 neural PCs (scaled for visualization). **I**. Example prediction of female walking speed from top PCs of a 20-neuron fast-adapt/slow-integrate MA population (averaged over 30 random instantiations of the population). **J**. Top: schematic showing song-evoked neural activity and walking speed regressor in neural PC space. Bottom: weights of each PC on the walking speed regressor, scaled by the square root of the total neural variance explained by each PC. **K**. Example 2-minute song segments driving neural activity either strongly (top) or weakly/negatively (bottom) for the top 4 neural PCs. **L-M**. Female walking speed variance explained from a 20-neuron fast-adapt/slow-integrate MA population either fixing *τ*_*a*_ and allowing *τ*_*int*_ to vary uniformly between 20 and 120 s (L) or fixing *τ*_*int*_ and allowing *τ*_*a*_ to vary uniformly between 100 ms and 2 s (M). (P < .005 between maximum walking speed variance explained with heterogeneous *τ*_*a*_ vs with heterogeneous *τ*_*int*_; 2-sided t-test).

To address the question of memory specifically, we asked which timescales within the MA responses were needed to predict locomotion. Resampling each model neuron’s *τ*_*int*_ and *τ*_*a*_ to take on either exclusively “fast” (100 ms - 2 s), “medium” (2 - 20 s), or “slow” (20 - 120 s) values across the population revealed the necessity of medium or slow integration and fast or medium adaptation. When adaptation was removed or integration timescales were fast, slow changes in walking speed were not well captured, whereas slow integration together with fast adaptation yielded predictions capturing both slow and fast changes in walking (Fig 2B-D). Of note, while medium integration and fast/medium integration could also explain a large amount of total walking speed variance (Fig 2A), this model yielded a poorer prediction of the low-frequency walking speed components (Fig S6A), a key component of female locomotion during courtship. Thus, the MA population’s ability to predict behavior by remembering song history is supported by slow integration dynamics and fast/medium adaptation. This reflects not all, but a sizeable fraction of the measured neural responses to sine/pulse block song stimuli (Fig S6B).

### The MA code for song history is multi-dimensional

How many model neurons and activity dimensions are needed by the “fast-adapt/slow-integrate” MA population? When we randomly generated variable-size MA populations in this regime, walking speed predictability plateaued near 15-20 model neurons (Fig 2E), suggesting the sufficiency a small collection of MA model neurons. As our model is noiseless, the improved prediction arises not through averaging but through heterogeneity of the population code. Curiously, although principal components (PC) analysis revealed the population responses to lie within a nearly 1-dimensional subspace, around 4-5 PCs were still needed to maximize walking speed predictions (Fig 2F-H), suggesting some low variance neural PCs in this code contain crucial behavioral information.

We found that the top PCs of these MA responses were highly interpretable. The top PC encoded the total amount of singing, the 2nd PC whether or not there was a recent period of relative quiet, the 3rd PC the sine-pulse ratio, and the 4th PC the mean duration of individual song modes (Fig 2J-K, S7-8). As we defined each PC to have a negative weight on the walking speed regressor (fit to the PC projections), an increase in the value of any of these features predicts female slowing, which may be a sign of “interest” in gathering more information from the male. Thus, the MA code we derived through NC automatically organizes interpretable song features along its top PCs. Extending previous work that predicted female slowing from bout duration and song amount (12), and sexual receptivity exclusively from pulse song (16), these results suggest that female responses to song are mediated by a rich space of sequential patterns, encoded in memory by multi-dimensional neural activity.

Finally, we asked whether heterogeneity in the adaptation vs integration time constants was more important. Using a population of 20 random MA neurons with either heterogeneous *τ*_*a*_ (uniform between 100 ms and 2s) and fixed *τ*_*int*_, or heterogeneous *τ*_*int*_ (uniform between 20-120s) and fixed *τ*_*a*_, we re-generated the artificial recordings to predict female walking speed. We found that fixing *τ*_*int*_ but retaining heterogeneous *τ*_*a*_ explained significantly more walking speed variance than the reverse (Fig 2L,M). Thus, heterogeneous adaptation dynamics are more fundamental to representing song history than heterogeneous integration timescales. This suggests a computational advantage of heterogeneous non-linear pre-processing of song through adaptation—increasing the dimensionality of the input to achieve richer sensitivity to recent temporal patterning—prior to integration.

In sum, female walking speed can be predicted from a neural memory of song history created as follows. Song is passed through a heterogeneous bank of nonlinear (multiplicative) adaptation processes, then integrated over slower timescales into persistent (slowly decaying) neural population activity. This produces a highly correlated but fundamentally multidimensional population code capable of accumulating song features over timescales up to minutes. We note that while NC has allowed us to *rule out* many encoding models, multiple solutions can still exist. For instance, one arrives at a slightly different MA population when building it up greedily (one neuron at a time), rather than randomly sampling from a fixed parameter regime, although the greedily built population is still organized around a backbone of heterogeneous MA responses with slow integration time constants (Fig S9). While in principle it may have been possible to identify this coding model and its influence on locomotion from head-fixed recordings in a rich virtual reality setup, the above validates NC as a drastically accelerated alternative process for generating quantitative evidence for and against different encoding models. Moreover, despite the clear limitation of only treating neural activity that can be predicted from behavioral observables, NC produces models directly refined on, hence immediately applicable to naturalistic behavior.

Having identified a candidate neural coding mechanism from the data through NC, we now turn to characterizing the coding dynamics and capabilities of MA model neuron responses to song sequences. In Fig 3 we investigate how single MA neurons respond to and compress song history. In Fig 4 we study the dynamics and computational advantages of a fast-adapt/slow-integrate population—although other response types (which in fact represent the majority of the whole-brain responses [Fig S6]) likely also play an important role in song coding, we focus our analysis on fast-adapt/slow-integrate neurons for the purposes of both simplicity and interpretability, and due to their central importance in predicting behavior.

**Fig. 3.**
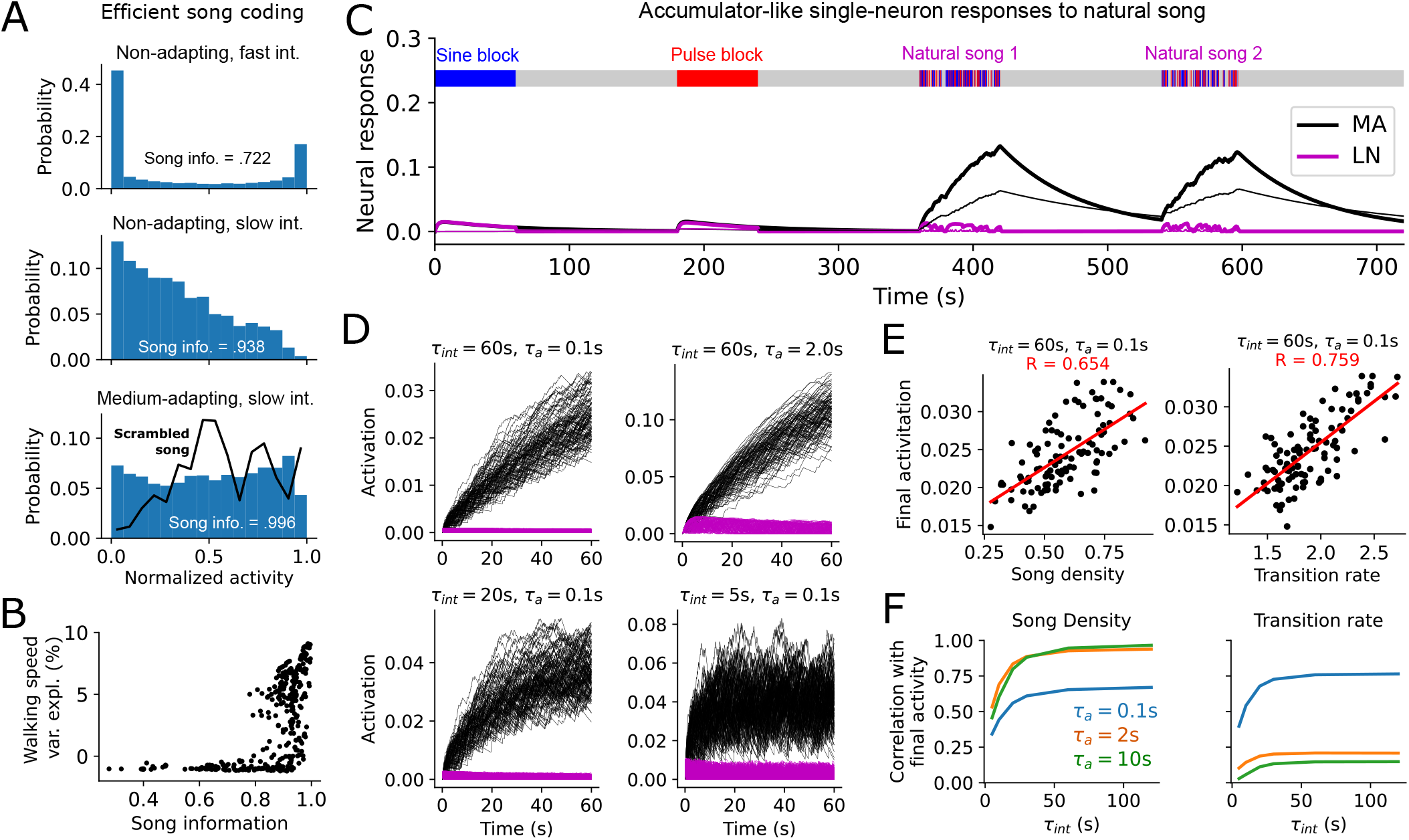
Compression and dynamical properties of single MA neurons in response to natural song. **A**. Histogram of normalized activity for three example neurons (top: *τ*_*int*_ = 0.5*s, τ*_*a*_ = ∞, *x*_*s*_ = *x*_*p*_ = 0.5, middle: *τ*_*int*_ = 120*s, τ*_*a*_ = ∞, *x*_*s*_ = *x*_*p*_ = 0.5, bottom: *τ*_*int*_ = 60*s, τ*_*a*_ = 10*s, x*_*s*_ = 1, *x*_*p*_ = 0) in response to 87 courtship songs (16 bins). Song information (given by response entropy [Eq. 4]) is reported relative to the entropy of a uniform distribution. Black line in bottom panel shows activity distribution in response to temporally scrambled songs (song information ≈ .943 for the scrambled song). **B**. Walking speed variance explained by single MA neuron responses to song vs song information retained in the MA neuron response (each point corresponds to one set of MA neuron parameters). **C**. Responses of 2 model MA neurons and their LN equivalents to a pure-sine block, a pure pulse-block, and 2 natural song segments. Thick trace: *τ*_*int*_ = 60*s, τ*_*a*_ = 2*s, x*_*s*_ = *x*_*p*_ = 0.5, thin trace: *τ*_*int*_ = 120*s, τ*_*a*_ = 0.5*s, x*_*s*_ = 0, *x*_*p*_ = 1. **D**. Responses of 4 example mixed-selectivity (*x*_*s*_ = *x*_*p*_ = 0.5) MA and equivalent LN neurons with different time constants to 108 different 1-minute song segments. **E**. Correlation of final neural activation after 1-minute natural song segment with two basic song features for the upper left neuron in D. Each black point corresponds to a different natural song segment. **F**. Correlations in E vs *τ*_*int*_ for three different values of *τ*_*a*_, with *x*_*s*_ = *x*_*p*_ = .5.

**Fig. 4.**
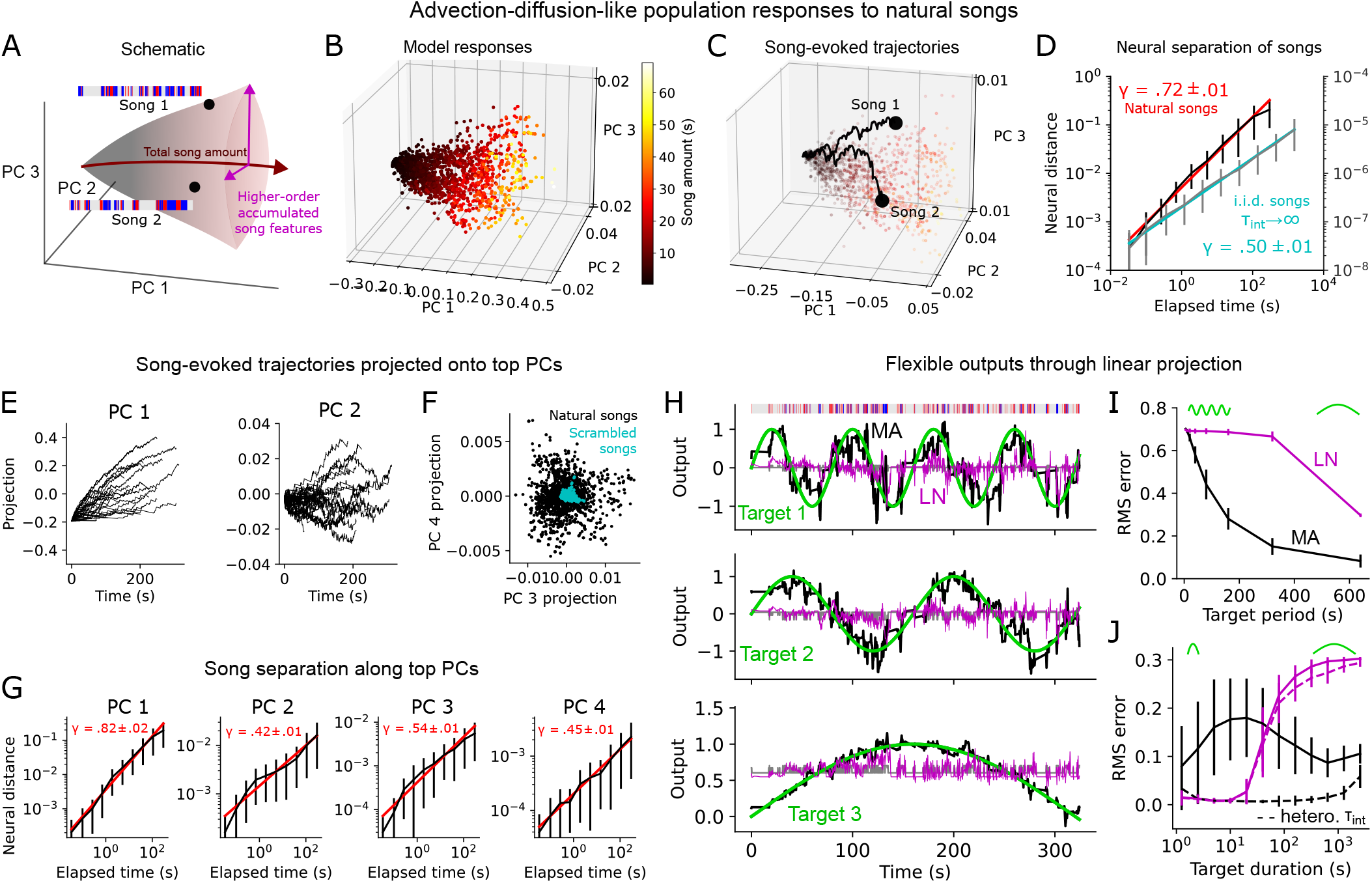
Advection-diffusion-like population trajectories as a general-purpose mnemonic representation. **A**. Schematic of population song representation from a fast-adapt/slow-integrate MA population. **B**. Responses of example 20-neuron MA population to randomly selected song segments spanning a range of durations, projected onto top 3 PCs of the song responses. In B-I parameters were set to be *τ*_*int*_ = 120s for all neurons, *τ*_*a*_ uniformly distributed between 100ms and 2s, and *x*_*s*_, *x*_*p*_ uniformly distributed between 0 and 1. **C**. Advection-diffusion-like population neural trajectories evoked by two example 30-second song segments. **D**. Euclidean distance in neural space between trajectories evoked by randomly selected 5-minute songs as a function of elapsed time (black). 57 5-minute songs were used (the full set was not used as many sessions ended before 5 minutes). Error bars show standard deviation across song pairs (100 pairs). Gray traces show same results for artificial i.i.d. songs 25 minutes long, using the same 20-neuron MA population except with *τ*_*int*_ = 10^6^ (with *τ*_*int*_ → ∞ representing an idealized version of this neural code). *γ* indicates mean and standard deviation of slope of best fit line on log-log plot (computed over 30 random selections of 100 song pairs). **E**. Example natural-song-evoked trajectories projected onto top PCs. **F**. Projections of song-evoked population responses (endpoints of trajectories) onto PCs 3 and 4, and comparison with scrambled songs. G. As in D but after projecting trajectories onto each of the top 4 PCs. **H**. Linear projections of song-evoked MA or equivalent LN population trajectories trained (Ridge Regression; *α* = 10^−15^) to reproduce three sine waves of varying frequency. Gray trace corresponds to target approximation directly from linear weighting of song inputs (binary sine and pulse inputs). **I**. Root-mean-squared error between target and prediction for 320s song segments at varying target periods using either MA or equivalent LN model. Error bars show standard deviation across 54 songs. **J**. As in I, except predicting targets from MA, LN trajectories generated by song segments of varying duration, with each target corresponding to a sine wave with half-period equal to the song segment duration. Dashed lines are populations with 20*s* ≤ *τ*_*int*_ ≤ 120*s*. Error bars as in I.

### Song history compression by MA neurons

We next sought to understand how the features of the MA neuron model shape the mnemonic encoding of song history. A natural question to ask is how much of the past is retained in the present—specifically, how much information about the history of song is stored in the momentary activity levels of our model neurons? To this end we asked how well a single MA neuron compresses song history into an instantaneous activity level *r*, quantified by the mutual information:

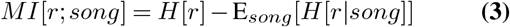

where *H*[*r*] and *H*[*r*|*song*] are the prior and conditional entropies of the activity of neuron (28). Because the MA neurons are deterministic, we have that *H*[*r*|*song*] = 0, hence

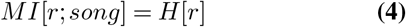

which can be estimated simply from the 1-D histogram of a model neuron’s activity *r* in response to the songs in the courtship dataset (aggregated over sessions and timepoints).

In general, different MA neurons had different response distributions, hence retained different levels of song information (Fig 3A, S10). For instance, model neurons with fast integration and no adaptation exhibited a bimodal activity distribution (Fig 3A, top), leading to a small entropy *H*[*r*] hence little song information. Model neurons with slower integration timescales yield a more heterogeneous activity distribution (Fig 3A, middle), as activity does not immediately decay during quiet periods. Model neurons with both slow integration and fast-to-medium adaptation exhibited activity distributions spread nearly uniformly across their total dynamic range (Fig 3A, bottom), suggesting these model neurons perform a near-optimal single-neuron compression of song. When these model neurons were presented with “scrambled” songs (breaking temporal correlations in song), their activity distribution became non-uniform (Fig 3A)–the high-entropy response distribution of these MA neurons is thus at least partly specific to naturalistic song, suggesting their response properties may be “matched” to song statistics. Overall, single MA neurons with high predictive power of behavior also contained high song information (Fig 3B)—in particular neurons with slow integration time constants (Fig S11). This suggests that these neurons are transmitting high-entropy song features, rather than arbitrary features buried in noise, and that they do so by remembering song history for long periods.

### Accumulator-like responses to natural song

To gain a deeper mechanistic understanding of how fast-adapt/slow-integrate MA neurons encode song history we re-examined the dynamics of their song-evoked responses. Reflecting our initial observations, the activity of these model neurons quickly plateaued in response to block sine or pulse song, due to the rapid adaptation (Fig 3C). The same neurons, however, produced a strikingly increased response to naturalistic song with a clear accumulator-like dynamic. This resembles neural correlates of evidence accumulation in other systems (29, 30), except here extending over minutes and crucially dependent on temporal patterns present in the song stimulus. Intuitively, the gaps and mode transitions in natural song allow adaptation to recover, with the slow integration allowing ongoing inputs to increment the current activity level.

Song accumulation dynamics occurred only in MA neurons and not in their LN equivalents (Fig 3C). Functionally, the initial nonlinearity of the MA neurons provides a means for distinguishing block from natural song in a manner that is not present in the LN neurons. Song-accumulation dynamics occurred for multiple MA parameter sets in the fast-adapt/slow-integrate regime, although the same song was represented slightly differently by different neurons. Variation of a single model neuron’s response to different songs typically increased over time (Fig 3D); and both song density and transition rates correlated with accumulation rate (Fig 3E), with stronger song-density correlations occurring for neurons with slower adaptation timescales and stronger transition-rate correlations for those with faster adaptation timescales (Fig 3F).

Thus, our model suggests that single neurons may encode naturalistic song history in memory via an accumulation process. Together with our information-theoretic analysis, this result suggests that adaptation and accumulation may interact with temporal statistics of natural social communication to efficiently transmit continuously unfolding information from the past into momentary neural activity levels.

### Advection-diffusion-like population dynamics

Next, we asked how the MA population model encodes the memory of song history collectively. To favor interpretability we examined a 20-neuron MA population with a fixed integration timescale *τ*_*int*_ = 120s and random fast adaptation timescales (100*ms* < *τ*_*a*_ < 2*s*), which accounted for most of the female locomotion variance explained by the original MA population (Fig 2M). Responses of this population to naturalistic song segments were confined to a cone-like region of the total neural response space, with the dominant axis (roughly the top PC) encoding total song amount and other dimensions higher order accumulated song features (Fig 4A-B, Fig 2K). Thus, the population responses to naturalistic song exhibit clear, multi-dimensional geometric structure.

Song-evoked neural population trajectories followed a striking advection-diffusion-like dynamic (Fig 4C). Classically, such dynamics describe the trajectory of a particle in a fluid flow subject to molecular or turbulent diffusion, serving, for instance, as an important model for odor plumes (31). Here the “advection” component, in analogy with downwind displacement, corresponds to the correlated accumulator-like response of all fast-adapt/slow-integrate neurons to song, reflecting the total amount of song heard. The diffusion component, arising in plumes from the accumulation of random collisions or turbulence, arises here from the highly fluctuating song inputs, with the heterogeneous adaptation dynamics transforming these into multi-dimensional signals that produce a Brownian-like dynamic upon integration. Note that the diffusion-like dynamics here are not noise but rather reflect deterministic transformations of song.

Functionally, these population dynamics continually separated neural representations of songs over a wide continuum of timescales. This means that early portions of song are not forgotten but rather contribute to the representation of song history up to minutes after they have passed. Indeed, the distance between song-evoked neural trajectories followed an approximate power law in time over nearly four decades (Fig 4D), with a scaling exponent of *γ* ≈ .72, suggesting a scale-invariant structure of the neural trajectories. Presenting scrambled songs with i.i.d. timepoints to the model also yielded advection-diffusion-like dynamics (with an exponent of *γ* ≈ .5, since we added no explicit heterogeneity across songs, which is likely present in the real data), indicating the scale invariance was not inherited from multi-scale song structure but is rather an emergent property of integrating adapted inputs. Multi-scale song separation occurred along many PCs of the population code (Fig 4E-G), recapitulating the multi-dimensional nature of these dynamics. Natural songs also evoked a larger response variance in higher PCs than scrambled i.i.d. songs (Fig 4F), suggesting that song temporal patterning may specifically engage MA neural dynamics to transmit more information into the female’s evolving neural code for song history.

Together, these results suggest (1) there exists a large, multi-dimensional space of persistent female neural states representing many different possible song histories (as opposed to, say, a few fixed-point attractors (32) visited in response to different song patterns); (2) the longer the song the more information is transmitted to the female, since representations of longer songs are further apart than shorter songs (a similar principle as has been observed in dynamic coding of olfactory stimuli, although over shorter timescales [seconds] (33, 34)); (3) the neural dynamics of encoding song history can in fact be partially understood as a variation of a well-known multi-scale physical process—advection-diffusion. Thus, the MA representation of song history not only improves behavioral predictions but reflects a rich, dynamic, information-dense, and interpretable mnemonic code.

### Flexible transformation of song trajectories

Finally, we asked whether the MA population coding algorithm, uncovered by studying song representations in the fly brain, could reflect a memory representation for general-purpose flexible computation. Despite the strong correlations, the multi-dimensional nature of the MA song code suggests that generic dynamical patterns might be extractable from the neural trajectory, similar to a reservoir computer—in which computations are performed via linear readouts of fixed random nonlinear transformations of time-series inputs (35, 36). To test this we trained linear readouts of the 20-dimensional neural trajectory evoked by a single example song segment to generate target output time-series with three different timescales (Fig 4H). We found that all three targets could be reproduced with reasonable accuracy from linear projections of the MA neural trajectory alone, but not from the matched LN trajectory, or from song itself. As all three targets are generated from the same song trajectory, a single song can hence be transformed via the MA code into a multi-dimensional behavioral signal, with different dimensions evolving at different timescales.

Across songs, the MA code outperformed the matched LN code over a range of target frequencies, although both failed at very high frequencies, likely due to the many periods present over a fixed-duration song (Fig 4I). However, faster target variations could be reproduced well by the MA code when short songs were used (Fig 4J). Further, when *τ*_*int*_ was allowed to be heterogeneous (20 < *τ*_*int*_ < 120s), rather than fixed, linear projections of the MA code could accurately reproduce the target sine wave output over several scales (Fig 4J). (Note that although a bank of non-adapting LN integrators can also be linearly transformed into very slow outputs, this occurs at the expense of being able to remember local temporal structure; Fig S12). Thus, the MA code reflects a reservoir-like memory mechanism that converts song history into multi-dimensional neural trajectories that can be linearly combined to produce flexible, scalable, multi-dimensional output signals, which could generically be used to drive complex behavior modulation.

## Discussion

We have shown how pure behavioral data can be used to test neural encoding models in naturalistic settings to identify novel coding principles underlying the neural basis of memory. Applied to fly acoustic communication (11, 12), our method revealed a candidate algorithm for creating and updating a continually growing representation of song history. Although the functional role of song memory in courtship remains unknown, our results that longer songs produce more unique (further apart) neural representations over multiple timescales suggests that longer songs transmit more information from the male into the female’s neural state, which would allow her to make a more informed mating decision.

Our work suggests a simple, general, and biologically plausible mnemonic coding algorithm—transforming input signals into slow, advection-diffusion-like neural trajectories via heterogeneous adaptation and slow integration—that could be applied in other systems that must process long, richly patterned input sequences (Fig 5). Such an advection-diffusion-like process also bears some resemblance to recent observations in mice performing a complex decision-making task, in which neural population activity traversed a nearly 1-dimensional “epoch”-ordered manifold (analogous to our first PC) in each trial, with trial-to-trial behavioral variability encoded in relatively small fluctuations within this manifold (37), producing new evidence that small neural fluctuations with rich dynamics may play an important role in processing information and guiding behavior.

**Fig. 5.**
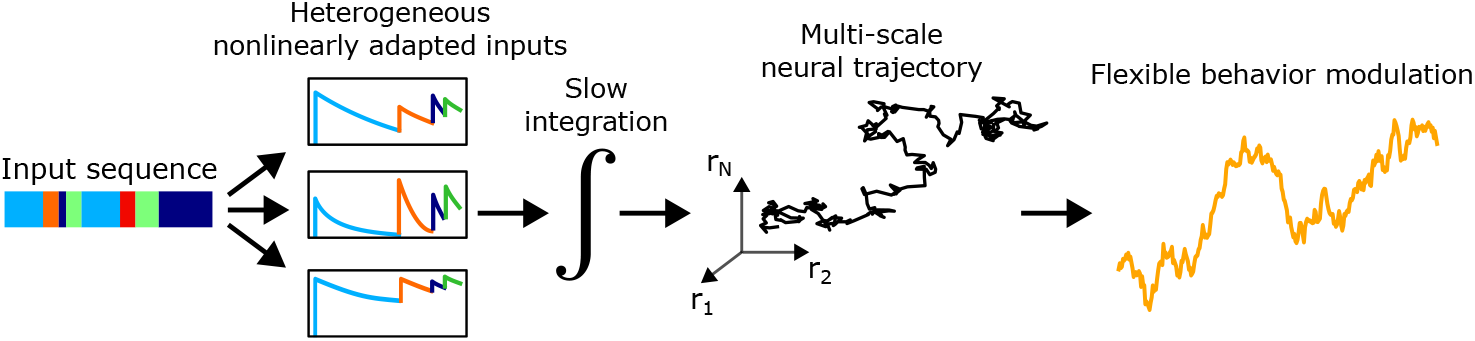
General neural principle for storing extended natural input sequences. Input sequences are passed through a bank of heterogeneous nonlinear adaptation processes, then integrated. This produces a multi-dimensional, multi-scale population trajectory that can be linearly projected to produce flexible signals to modulate behavior.

### Generality and limitations of Natural Continuation

NC is an approach for quantitatively evaluating neural coding models using unrestrained behavior data. This serves as a powerful complement to methods for decomposing naturalistic behavior into motifs and their transitions in order to guide subsequent neural experiments (38–42). By capitalizing on naturalistic behavior, NC overcomes restrictions imposed by limited neural recordings, providing a widely applicable approach for extending insights from the existing body of neural data and models to the rapidly growing collection of naturalistic behavioral data. One exciting direction, for instance, would be to use NC to resolve dynamic neural computations during naturalistic odor-tracking, by combining the rich literature on olfactory neural responses (13, 25, 34, 43, 44) with behavioral recordings of flies following turbulent plumes (45, 46).

Nevertheless, NC’s ability to augment discrimination power over neural encoding models is subject to important limitations. First, the behavior of interest must be present and resolvable in the behavioral data. Second, mechanistic insight is limited by the set of neural encoding models selected. Third, all results are mediated by the encompassing behavioral model in which the hypothesized neural dynamics are embedded. In general, both the neural and behavioral models should be chosen in accordance with the question to be investigated, i.e. with a “scientist-in-the-loop”. Finally, even given a rich behavioral time-series, multiple neural models may predict the behavioral data equally well—ultimately, the power of NC is instead to rule out possible models, in order to guide the search for neurally plausible solutions to complex problems faced in natural settings.

### Process memory

The MA model for song history we have studied here falls within the framework of *process memory* (47). In contrast to searching for specific neural circuits that allow animals to maintain stimulus information through a delay period (48–50), process memory posits that most neural circuits have intrinsic mnemonic capabilities that are continually engaged and fundamentally entangled with online information processing. While better suited for studying memory-dependent behaviors without explicit storage, maintenance, and retrieval periods, the lack of separation between memory and information processing in this framework introduces substantial complexity. Our analysis of fly courtship, however, reveals that process memory in this and potentially other systems may be supported by relatively simple mechanisms that nonetheless have rich mnemonic capabilities.

### Connection to diffusion models in neuroscience and machine learning

The most well-studied role of diffusion dynamics in neuroscience is in drift-diffusion models (DDM) of evidence accumulation (29, 51). While our model shares the core principle of integrating its inputs, it differs in three key ways: (1) it takes place over a much longer timescale (minutes) then typical DDM experiments (seconds); (2) instead of directly receiving two competing input streams, the model converts a single ternary input stream into a multi-dimensional format via the bank of adaptation processes (note also that each adaptation variable does not encodes a specific “feature” of song—rather the heterogeneous population collectively encodes something closer to a multi-dimensional continuum of song features); (3) our model here is noiseless, hence the diffusion dynamics are pure signal, unlike classic DDM in which much of the diffusion is due to noise. In our model such diffusion-like encoding leads to multi-scale separation of songs and produces a neural trajectory that can be linearly transformed into flexible multi-scale outputs.

The diffusion-like component of our model may in fact be more similar to diffusion models of image generation in machine learning (52, 53), where diffusion-like dynamics are key to generating meaningful outputs. Ongoing efforts have sought to apply the benefits of diffusion to tokenized language processing (54), which is currently dominated by transformer architectures (55), with most approaches focusing on iteratively refining an entire text. Our work suggests a different relationship between diffusion and tokenized inputs in the fly brain: the sequential presentation of token-like inputs (song elements) deterministically drives a diffusion-like neural trajectory to encode the input history. In fact, the MA population model is strikingly similar to a recently explored simple but powerful scheme for sentence embeddings, in which the constituent word embeddings (analogous to our nonlinear transformation of recent song patterning) are simply weighted and summed to represent the sentence (56). Our model can also be interpreted as a locality-sensitive hashing operation (57) for transforming variable-length input histories into advantageous fixed-dimensional vector codes, in our case retaining rich mnemonic information. Incorporating such a process into a language model may provide a simple means for representing unbounded input histories.

### Network and cellular mechanisms

We have focused on neural *encoding*, building on a long line of sensory neuroscience research (23, 30, 58–60). Neural responses to song, however, emerge at a mechanistic level from physical network and/or cellular processes (18, 61, 62). Our work points to the necessity of adaptation and integration, which could each arise via either network or cellular mechanisms. For instance, although identifying robust network mechanisms for integration remains an open challenge in neuroscience (63), inputs may also get integrated by intracellular mechanisms such as calcium-sensitive cation currents (64, 65) or intrinsic excitability changes (66), which would make the mechanism robust against network variability and obviate any need for fine-tuning connectivity. Adaptation could also arise either at a network level or locally; the fact that adaptation parameters are generated randomly in our model suggests this process would also be robust to network variability. In general, understanding network interactions in this system would also help constrain models for spontaneous activity variability unrelated to song history, refining predictions of response variability over time or individuals. Although we focused on memory via persistent neural activity, our results also do not rule out the role of plasticity in the preservation of song memories beyond the minutes-long timescale. Plasticity in the mushroom body, a canonical insect memory center (67, 68) that also exhibits auditory responses (17), in principle could reformat song representations into a synaptic code to persist information after activity decays.

### Role of other song-responsive neurons

Our work focused on fast-adapt/slow-integrating MA neurons, but neural responses with faster integration time constants are clearly visible in the head-fixed recordings (Fig 1E, S1) (17, 18). While we found that these features were not needed to predict female walking speed in the courtship dataset we studied, they may be relevant for other purposes, for instance integrating song memory with ongoing multi-sensory information, mediating vaginal plate opening or ovipositor extrusion (16, 69) or responding to interactions between competing males (70). Functionally, shorter-timescale responses, along with neurons exhibiting sine-offset responses (Fig S3), may also help extend neural dynamics through quiet periods in song. These could then be used to generate song-dependent dynamical behavioral modulation that continues even when singing has ceased. Shorter timescale song responses may also reflect network processes used to form the fast-adapt/slow-integrate responses.

### Importance of song structure

Our work reveals several functionally relevant ways in which song temporal structure may interact with the dynamical responses of fly auditory neurons. First, gaps and transitions in song allow the adaptation variables of the model to recover, enabling naturalistic song to drive model neural responses much more strongly than single-mode song blocks (Fig 3C), in which adaptation causes responses to quickly plateau. Second, naturalistic temporal correlations cause behaviorally predictive MA neurons to more evenly explore their full dynamic range, suggesting an efficient single-neuron code for song history (Fig 3A). Third, naturalistic song structure drove increased exploration along the 3rd and 4th PCs of the population neural code over scrambled song, i.e. “spreading out” the representation (Fig 4F). Although adaptation has long been theorized to enhance neural coding by modulating responses to changing stimulus statistics (71–75), here we have shown how adaptation can support the efficient encoding of the stimulus temporal structure itself, potentially serving to efficiently encode an entire input history. Our work also further refines a hypothesis for the functional role of the complex temporal structures of courtship song: whereas a white-noise-like song could in principle carry more information than structured song (28), if song must be processed by a neural dynamical system limited by biophysical responses, one will generally expect the optimal information-transmitting song to have more structure than white noise, which may be the case in fly courtship.

### Predictions

Our model yields testable predictions about neural coding during both fly social communication and memory-dependent behaviors in other animals. First, a population of neurons in the female fly brain should exhibit an increased response to natural over block song, since the gaps and transitions in natural song allow the adaptation variables mediating song responses to recover. Second, single neurons in the fly brain, or potentially single dimensions of the population response, should exhibit an accumulator-like dynamic in response to natural song that roughly encodes how much total song has transpired. Third, typical pairs of natural songs should become more discriminable in their population code over multiple timescales (Fig 4D). Fourth, natural songs, and potentially other natural inputs that must be integrated over long periods, should evoke multi-scale advection-diffusion-like dynamics in the brain. We additionally predict that such multi-scale dynamics may be a generic neural correlate of fluctuating language-like inputs that must be remembered over long timescales, e.g. in acoustic communication in mice (76), or narrative processing in humans or language models (2). This could be used, for instance, to inform the search for neuroimaging biomarkers for memory disorders (77). We note that in general we may also expect these dynamics to be multiplexed with other neural processes.

### Origin of multi-scale behaviour

A pervasive characteristic and challenge of naturalistic animal behavior is that it unfolds over multiple timescales (39, 78, 79). How these timescales are organized and emerge from the many components of the nervous system and the animal’s interaction with its environment remains a basic open challenge. One central hypothesis is that multi-timescale neural dynamics, which could in turn drive multi-timescale behavior, emerge from a hierarchy of brain areas with different recurrent processing dynamics (47, 80). Our model, however, suggests that multiple timescales can also emerge from a distinct, complementary mechanism: nonlinear expansion and transformation of a stream of rapidly fluctuating inputs, followed by integration, producing a multi-dimensional Brownian-like trajectory. This represents an alternative source of multi-scale neural dynamics, with rich mnemonic structure, which could be engaged for general behavioral modulation or potentially for complex tasks such as language processing.

## Methods

### Neural and behavioral data

We used 224 calcium recordings (GCaMP6s indicator, sampled at 8Hz) distributed across 50 different cell lines (multiple flies per cell line) in response to 10-second blocks of pure sine or pure pulse song presented through a speaker. Details can be found in (18). Responses analyzed in Fig S1B,D and S6B were from panneuronal imaging experiments in (17). We note that although GCaMP6s has a time constant of ∼ 1s, in contrast to faster indicators, it is unlikely that this influenced our findings because (1) the integration timescales we have studied are much longer (up to minutes), and (2) NC enables the general comparison of encoding models informed by neural data, which would rule out any encoding models fit to possibly confounded neural data if they don’t generalize to predict behavior. Courtship sessions were collected and described in detail in (11). Females were blind and pheromone-insensitive to increase their auditory responses; otherwise both flies were free to court in the arena. Songs were recorded with microphones on the arena floor, then segmented into sine/pulse timepoints; locomotion was recorded via overhead cameras then manually/automatically tracked.

### Neural encoding models

For each neuron we fit the MA model using gradient descent over its 4 parameters: *τ*_*int*_, *τ*_*a*_, *x*_*s*_, *x*_*p*_, minimizing the squared error between the model’s response and the trial-averaged calcium responses to the sine and pulse blocks (including the 10-second stimulus and a 10-second post-stimulus period). To fit the LN model we derived an analytical relationship allowing us to parameterize the filters *h*_*s*_, *h*_*p*_ by the same 4 parameters as the MA model, such that the LN and MA model had the same step response to sine and pulse input. We then performed gradient descent over these 4 parameters to adjust the filters to capture the offset response as well. The nonlinearity was signed rectification (based on the sign of the calcium responses). We fit linear readouts from song-evoked activity in the courtship sessions to female walking speed using Ridge Regression.

Further methods details can be found in the Supplement.

## Acknowledgments

We thank William Bialek, Albert Lin, Wayan Gauthey, Nicolas Lenner, Luca di Carlo, Daniel Weilandt, Kevin Chen, Agostina Palmigiano, as well as members of the Pillow, Murthy, and Bialek groups for many helpful discussions regarding the development of this project.

## Funding

This work was supported in part by a Simons Collaboration for the Global Brain Investigator Award (to MM and JP); the National Institutes for Health BRAIN Initiative (1R01NS104899-01) (to MM and JP); the Swartz Theoretical Neuroscience Fund (to RP); the Jane Coffin Childs Memorial Fund for Medical Research (to CB).

## Notes

http://arks.princeton.edu/ark:/88435/dsp01rv042w888

http://arks.princeton.edu/ark:/88435/dsp016395wb26g

http://arks.princeton.edu/ark:/88435/dsp01h702q942c

### Competing Interest Statement

The authors have declared no competing interest.

