## Supplementary Information for "Inferring neural dynamics of memory during naturalistic social communication"

**This PDF file includes:**

SI References

### Extended methods

**Neural data.** We fit our encoding models to 224 neural recordings obtained via calcium imaging, the majority of which were recently published in (1). Specific cell types were targeted using sparse, specific split-GAL4 lines and the fluorescent calcium indicator GCaMP6s. Imaging was performed one line at a time, i.e. only a single cell type was imaged in each experiment. Stimuli were presented in a block-randomized order with 10 s sine and pulse blocks (as well as a 10 s white noise block) interleaved with 20 s inter-stimulus intervals. All stimuli were presented through a speaker to a female fly head-fixed under a two-photon calcium imaging microscope. Fluorescence changes were then extracted from the resulting video. Only responses that were statistically significantly time-locked to stimulus onset were kept. See (1) for further details. Occasionally multiple regions-of-interest (ROI) were identified in a single cell line and treated separately. Moreover, sometimes the same cell line was imaged in multiple flies. In our analyses we treat all ROIs across all experiments separately—for simplicity, we have referred to each ROI in each experiment as a "neural recording". For each neural recording, responses were averaged across trials, then z-scored across all timepoints. In our analyses here we have only used the stimulus periods encompassing the 10-second pure sine or pulse blocks and their surrounding pre- and post-stimulus periods.

The neural responses analyzed in Fig S1B,D and Fig S6B were from (2). These were collected in a similar manner as those in (1), except using pan-neuronal imaging instead of individual lines for different cell types, and with 10-s stimulus blocks presented in a stereotyped order (pulse, sine, white-noise). See (2) for details.

**Behavioral data.** We used publicly available naturalistic courtship data described in (3–5). In each session/ritual a virgin male and female fly were placed in the arena and allowed to interact for up to 30 minutes. If copulation occurred before 30 minutes had passed the session was ended at the time of copulation. Locomotion was recorded via overhead video cameras and song through floor microphones, and subsequently processed using a combination of automated and manual segmentation and tracking. Both locomotion and the ternary (quiet, sine, pulse) song time-series were sampled at 30.03 Hz (the auditory recordings of song were initially conducted at a much higher sampling rate, then segmented and binned into 30.03 Hz sampling bins, which was the sampling rate of the video cameras used to record locomotion). Female flies were also rendered blind and pheromone insensitive (PIBL) to increase their responses to auditory stimuli. See (3) for further details on the data collection.

For the analyses reported in this manuscript we used 87 courtship rituals (comprising 13.4 hours of song/behavior) with wild type PIBL females and two strains of males (NM91 and ZH23), which sang more robustly than other male strains and evoked robust female locomotion. Although more courtship rituals were available, we found that including them in our analysis did not substantially improve our MA-based predictions of female behavior; and the improved predictive capacity using the MA over the LN model remained as more trials were included in the training data (Fig S13). Using 87 sessions also greatly simplified the computational resources required to generate and store the many iterations of artificial neural recordings and behavioral predictions that we performed.

**Fitting the encoding models.** We fit the four parameters of the MA encoding model to each neuron by minimizing the squared error between the model MA response to the sine and pulse stimuli (together with their 10-second post-stimulus periods) and the empirical calcium responses. To ensure fair comparison between the abilities of the LN vs MA models, we derived a method to parameterize the LN models by exactly the same parameters as the MA model ($\tau_{int}, \tau_a, x_s, x_p$) (otherwise the LN model would have many more parameters, including each timepoint in each filter). To this end, we analytically computed the MA step response and then took its time-derivative to identify the corresponding LN filter, which was consequently parameterized by the MA parameters (see next section). For the nonlinearity $g$ we used a signed rectifying nonlinearity in accordance with whether the empirical neural activity increased or decreased following stimulus onset. We then fine-tuned the LN filter by adjusting the four parameters through gradient descent so that the LN response captured both the onset and offset responses of the block stimuli.

We also fit the LN model directly to the calcium data using Ridge Regression and a sigmoidal nonlinearity. The nonlinearity was

$$g(z) = r_{min} + (r_{max} - r_{min}) \left[ \frac{\tanh\left(\beta(z - z_0)]\right) + 1}{2} \right], \quad [1]$$

where $z$ is the filtered song input ($h_s * I_s + h_p * h_s$). We fit the filters first without the nonlinearity, then fit the nonlinearity given the filters, and finally performed a joint optimization of both the filter and nonlinearity parameters to fine tune the fit, minimizing the squared error between the LN predictions and the mean calcium responses, and penalizing the filter weights with the same Ridge Regression parameter. We also found that using a sigmoidal nonlinearity produced similar results as a signed rectification nonlinearity. In general, however, the preceding fitting procedure in which we parameterized the LN model by the MA parameters yielded more conservative results (i.e. the LN models parameterized by the 4 MA parameters yielded a better LN prediction of female walking than the LN models fit with Ridge Regression), in particular since the LN model tended to inherit very similar timescales as the MA timescales (Fig S15).

**Procedure for constructing LN encoding model parameterized by MA parameters.** First we derive an analytical form of the step response of the MA neuron. For $I_s = \Theta(t), I_p = 0$, where $\Theta$ is the Heaviside step function, we have

$$a_s(t) = (1 - \exp(-t/\tau_a))\Theta(t) \quad [2]$$

hence

$$\tau_{int}\frac{dr}{dt} = -r + x_s \exp(-t/\tau_a)\Theta(t). \quad [3]$$

Then we can write

$$r_s^{step}(t) = h_r * x_s \exp(-t/\tau_a)\Theta(t) \quad [4]$$

where

$$h_r(t) = \frac{1}{\tau_{int}} \exp(-t/\tau_{int})\Theta(t) \quad [5]$$

is the exponential filter described by the dynamical system for $r$. That is,

$$r_s^{step}(t) = \int_0^\infty h(t')u(t-t')dt' = \frac{x_s}{\tau_{int}} \int_0^\infty \exp(-t'/\tau_{int}) \exp(-(t-t')/\tau_a)\Theta(t-t')dt' \quad [6]$$

where the $\Theta(t)$ corresponding to the filter $h_r$ has been absorbed into the limited integration range. We can take care of the second heaviside function in the same way:

$$r_s^{step}(t) = \frac{x_s}{\tau_{int}} \int_0^t \exp(-t'/\tau_{int}) \exp(-(t-t')/\tau_a)dt' = \frac{x_s}{\tau_{int}} \exp(-t/\tau_a) \int_0^t \exp(-t'/\tau_{int}) \exp(t'/\tau_a)dt'. \quad [7]$$

When $\tau_{int} = \tau_a$ we have

$$r_s^{step}(t) = \frac{x_s}{\tau_a} t \exp(-t/\tau_a) \quad [8]$$

i.e. an alpha function since the integral becomes $t$. Otherwise

$$\begin{aligned} r_s^{step}(t) &= \frac{x_s}{\tau_{int}} \exp(-t/\tau_a) \int_0^t \exp\left(-t'(1/\tau_{int} - 1/\tau_a)\right) dt' \\ &= \frac{x_s}{\tau_{int}} \exp(-t/\tau_a) \frac{\exp\left(-t'(1/\tau_{int} - 1/\tau_a)\right)}{1/\tau_a - 1/\tau_{int}} \Bigg|_0^t \\ &= \frac{x_s}{\tau_{int}} \exp(-t/\tau_a) \frac{\left(\exp\left(-t(1/\tau_{int} - 1/\tau_a)\right) - 1\right)}{1/\tau_a - 1/\tau_{int}} \end{aligned} \quad [9]$$

Simplifying, we have

$$r_s^{step}(t) = \frac{x_s}{\tau_{int}/\tau_a - 1} \left(\exp(-t/\tau_{int}) - \exp(-t/\tau_a)\right) \quad [10]$$

i.e. the scaled difference between two exponential filters with timescales $\tau_{int}$ and $\tau_a$.

The sine filter is given by the derivative of the step response. When $\tau_{int} = \tau_a$ we have

$$h_s(t) = \frac{d}{dt} r_s^{step}(t) = \frac{x_s}{\tau_{int}} \left(\exp(-t/\tau_a) - \frac{t}{\tau_a} \exp(-t/\tau_a)\right). \quad [11]$$

Otherwise

$$h_s(t) = \frac{x_s}{\tau_{int}/\tau_a - 1} \left(\frac{-1}{\tau_{int}} \exp(-t/\tau_{int}) + \frac{1}{\tau_a} \exp(-t/\tau_a)\right) \quad [12]$$

And similarly for the pulse filter:

$$h_p(t) = \frac{x_p}{\tau_{int}} \left(\exp(-t/\tau_a) - \frac{t}{\tau_a} \exp(-t/\tau_a)\right) \quad [13]$$

when $\tau_{int} = \tau_a$ and

$$h_p(t) = \frac{x_p}{\tau_{int}/\tau_a - 1} \left(\frac{-1}{\tau_{int}} \exp(-t/\tau_{int}) + \frac{1}{\tau_a} \exp(-t/\tau_a)\right) \quad [14]$$

otherwise. Note that the $\tau_a \neq \tau_{int}$ case converges to the $\tau_a = \tau_{int}$ case as $\tau_a \to \tau_{int}$, even though the denominator in the prefactor goes to 0.

To construct the LN model we use these filters, combined with a signed rectification nonlinearity in accordances with the signs of the selectivities $x_s$ and $x_p$. Thus, by construction the LN responses to the step inputs i.e. block song onset are identical to the MA responses. However, the offset responses following the 10-second block stimulus, while typically similar, will not necessarily be exactly matched to the MA responses, however. Therefore we adjusted the 4 parameters of the LN model using a standard gradient-descent procedure (scikit-learn's minimize function) to maximally reproduce the full block song responses over the 10-second stimulus period and 10-second post-stimulus period.

**Bout-duration and hand-picked feature models.** It was shown in Clemens, et al (2015) (4) that among several manually chosen song features, time-averaged bout-duration exhibited the strongest correlation with time-averaged female walking speed, with the correlation plateauing near a 1-minute averaging window. To verify these conclusions within our moment-to-moment predictive analysis of female walking speed, we computed the momentary values of a battery of hand-picked song features, time-averaged over different windows, and then used each feature individually to predict moment-to-moment female walking speed (using either a 1-second or 1-minute forward averaging window for walking speed, as in our other analysis). Corroborating the results of Clemens et al we found mean bout duration, averaged over about 1-4 minutes, provided the strongest prediction of moment-to-moment female walking speed, explaining about 10-12% of the variance of the 1-second-averaged walking speed and 12-15% of the variance of the 1-minute-averaged walking speed (Fig S14). (As in our other analyses, the bout-duration regressor was fit on 80% of the trials and the prediction and variance explained computed on a test/held-out 20% of the trials, and finally averaged over 30 training/test splits). Note that while (4) showed that mean bout duration could be estimated with a single-neuron filter applied directly to the envelope of song acoustic envelope waveform, combined with a set of nonlinearities and an integrator, we did not explicitly implement this pseudo-neural model, since its end result to was to provide a highly accurate estimate of mean bout duration (Pearson correlation = 0.93), which we instead computed directly from the ternary song representation.

**Direct song-to-locomotion filter.** To estimate how much female locomotion variance could be explained by direct linear filters on sine and pulse song, we represented both a sine and pulse filter, $h_s$ and $h_p$ as a sum of 16 raised cosine basis functions:

$$b_i(t) = \frac{1}{2}\cos(a\log(t+c) - \phi_i) + \frac{1}{2} \qquad [15]$$

where we set $a = 6$, $c = 0$, and $\phi_i = i\pi/2$, which spanned timescales up to approximately 1 minute. Filters were then represented as

$$h_s(t) = \sum_{i=0}^{15} w_s^i b_i(t) \qquad [16]$$

and similarly for the pulse filter. We then fit the basis function weights $\{w_s^i\}, \{w_p^i\}$ using Ridge Regression ($\alpha = 10$) to minimize the squared error between female locomotion and the summed filter outputs (Fig S17).

**Fitting the neural-to-locomotion readout.** Except when otherwise mentioned, each encoding model and variation we investigated produced a 224-dimensional time-series (the number of original calcium recordings) accompanying each of the 87 courtship sessions. Each recording was a deterministic function of the courtship songs, as all encoding models were deterministic. As our locomotion variable we used the total walking speed of the fly, which could comprise both forward and lateral components (relative to the fly's body axis), and which was as or more predictable than purely forward or purely lateral motion (Fig S4).

To predict locomotion we first "forward-smoothed" the locomotion time-series (i.e. total walking speed at any timestep) by replacing the walking speed at each timestep $t$ with its time average from $t$ to $t + \Delta T$. Our motivation for smoothing only in the forward direction was to not contaminate our measure of ongoing/future locomotion with past features of female locomotion that could have influenced song at time $t$.

For each encoding model or variation we trained a linear readout from the artificial population neural activity at time $t$ (which encodes song history up till time $t$) to the forward-smoothed female locomotion variable at time $t$. We trained the readout (using Ridge Regression with $\alpha = 10$) on 80% of the courtship sessions and tested the readout across all timepoints in the remaining 20% of courtship sessions, which contained different flies. In general, female flies walked even before male song began. To focus on song-modulated locomotion, we therefore excluded from training/testing (1) all timepoints occurring before the first non-quiet song timepoint, (2) any timepoint occurring after more than 30 seconds of pure quiet had passed. (Note that this second condition was extremely rare, since the mean quiet period was only 2.34 seconds [Fig S4], with substantially longer quiet periods occurring with very low probability.) To compute the final score for each encoding model we performed this procedure for 30 random 80/20 splits of the 87 sessions into training/test sessions and reported the variance explained in held-out sessions, averaged across splits.

**Song shuffling procedure.** Songs were shuffled (Fig 1J) as follows. Songs could not be randomly assigned to different courtship sessions directly because sessions had different lengths. Instead, we concatenated all songs, then randomly circularly shifted them, then re-segmented the songs into individual sessions using the empirical session durations. This procedure retains temporal correlations in song while breaking correlations between song and locomotion.

**Estimating song information.** For a given MA model neuron we estimated the information about preceding song it contained in its instantaneous activity level by presenting the entire courtship song extracted in each of the 87 sessions to the neuron (excluding any initial quiet period at the start of a session before singing began), and then creating a histogram of responses (using 16 evenly spaced bins ranging from 0 to the maximum neural response level) aggregating over all songs and timepoints (yielding 1,448,116 timepoints total binned into 16 bins). We then computed the Shannon entropy of the resulting histogram:

$$H[r] \equiv -\sum_i f_i \log f_i \qquad [17]$$

where $f_i$ are the estimated fraction of timepoints in each bin. While this quantity is in general a biased estimate of entropy, we do not expect our results to be significantly affected, since the histogram estimate is built from over 1 million timepoints. Values reported in the figures are relative to the Shannon entropy of the equivalent uniform distribution, hence range between 0 and 1.

**Stimulus-invariant adaptation model.** The stimulus-invariant version of the MA encoding model was given by

$$\begin{aligned} \tau_{int}\frac{dr}{dt} &= -r + x_s(1-a)I_s(t) + x_p(1-a)I_p(t) \\ \tau_a\frac{da}{dt} &= -a + I_s(t) + I_p(t) \end{aligned} \quad [18]$$

i.e. reproducing the original MA model except with only a single adaptation variable $a$.

**Greedily constructed behaviorally predictive MA population.** Our greedily constructed behaviorally predictive MA population was built in the following way. We first defined a finite set of parameter values to select from, since this procedure amounts to an optimization requiring training a readout and predicting locomotion over 30 training/test splits of a large behavioral dataset for every iteration, making it infeasible to easily search over a continuous parameter space. The range of parameters we considered was $\tau_{int} \in \{0.1, 0.5, 1, 2, 5, 10, 30, 60, 120\}$ s, $\tau_a \in \{0.1, 0.5, 1, 2, 5, 10, 30, 60, \infty\}$ s, $x_s \in [0, 0.5, 1]$, $x_p = 1 - x_s$. Note that we did not include negative selectivities, which correspond to empirical neural activity that decreases in response to song, since this does not affect the ability to predict behavior due to the ability of the linear readout to absorb arbitrary signs and scalings of the neural activity.

We first generated an artificial recording containing one neural response for every combination of parameters. Next, we identified the single neuron that could best predict female locomotion in held-out courtship sessions. We then iterated over selecting the next best neuron, that if added to the existing population, maximally increased female locomotion variance explained in held-out courtship sessions, averaged over training/test splits, up to 50 neurons.

**Selection of songs for accumulation and manifold analysis.** To investigate accumulator-like dynamics of the MA neurons we presented 1-minute song segments taken from the courtship rituals used in the rest of our analyses. To extract song segments, we segmented all songs across all 87 sessions into song segments lasting at least 1 minute and separated by at least 5 seconds of quiet. For any song segments selected as such that extended beyond 1 minute, only the first minute was used. This yielded 108 unique natural song segments, which we presented to the model neurons studied in Fig 3.

For the manifold analysis we curated a collection of song segments of 40 different durations $T$ ranging from 1s to 300s, spaced either linearly (Fig 4B-C) or logarithmically (Fig 4D,G). To extract segments of length $T$, each courtship session was segmented into segments of duration $T$ s or greater separated by at least $T$ s of no singing, then truncated to have length exactly $T$. Principal component analysis in Fig 4E-G was performed on the responses of the fast-adapt-slow-integrate 20-neuron MA population to these song segments.

**Encoding models reproducing offset responses.** To allow our encoding models to reproduce the occasional sine-offset responses seen in the calcium recordings in (1) (Fig S3), we used an MA model in which quiet periods during song were treated as their own song mode (excluding the initial quiet period at the beginning of each session), which allowed us to simply augment the MA model with a third selectivity $x_q$ and adaptation variable $a_q$ that had the same structure the sine and pulse adaptation selectivities and adaptation variables (Fig S17). We also considered an LN model, named "LN-ReLu-Flex" (Fig S17), in which the nonlinearity was a piecewise linear function (with one hinge at the origin) that was not constrained to be strictly monotonic.

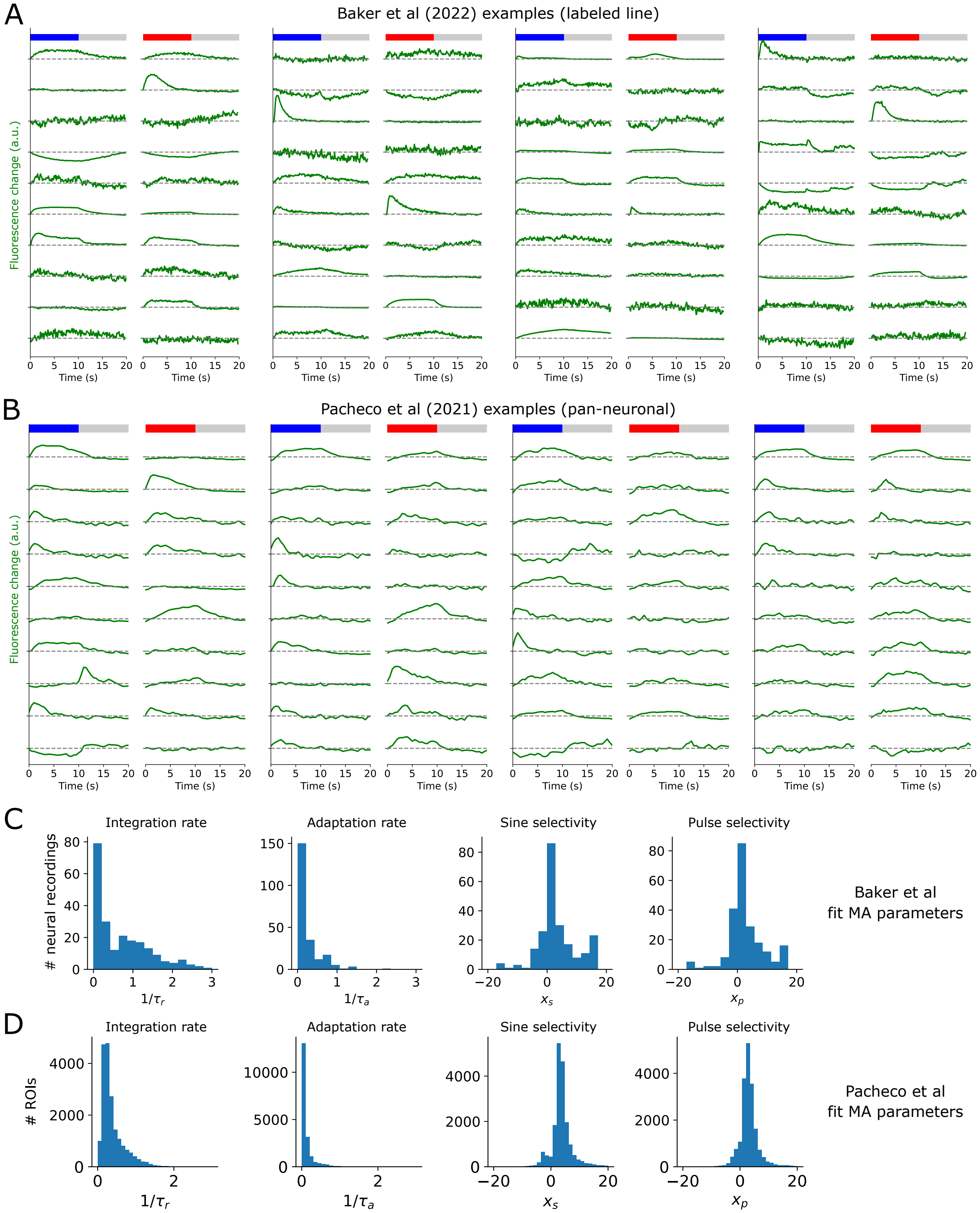


**Fig S1. Example labeled line and pan-neuronally imaged calcium response to block song.** A. Example responses from a diverse selection of neurons in the labeled line data (Baker et al 2022) (1). Every pair of responses to the sine and pulse blocks corresponds to one neural recording. B. Example responses from a selection of example ROIs extracted from the pan-neuronal imaging data in Pacheco et al. 2021. C. Distributions of MA parameters fit to all neurons in Baker et al 2022 dataset (N=224). D. Distributions of MA parameters fit to all ROIs in Pacheco et al 2021 dataset (N=19036) (2).

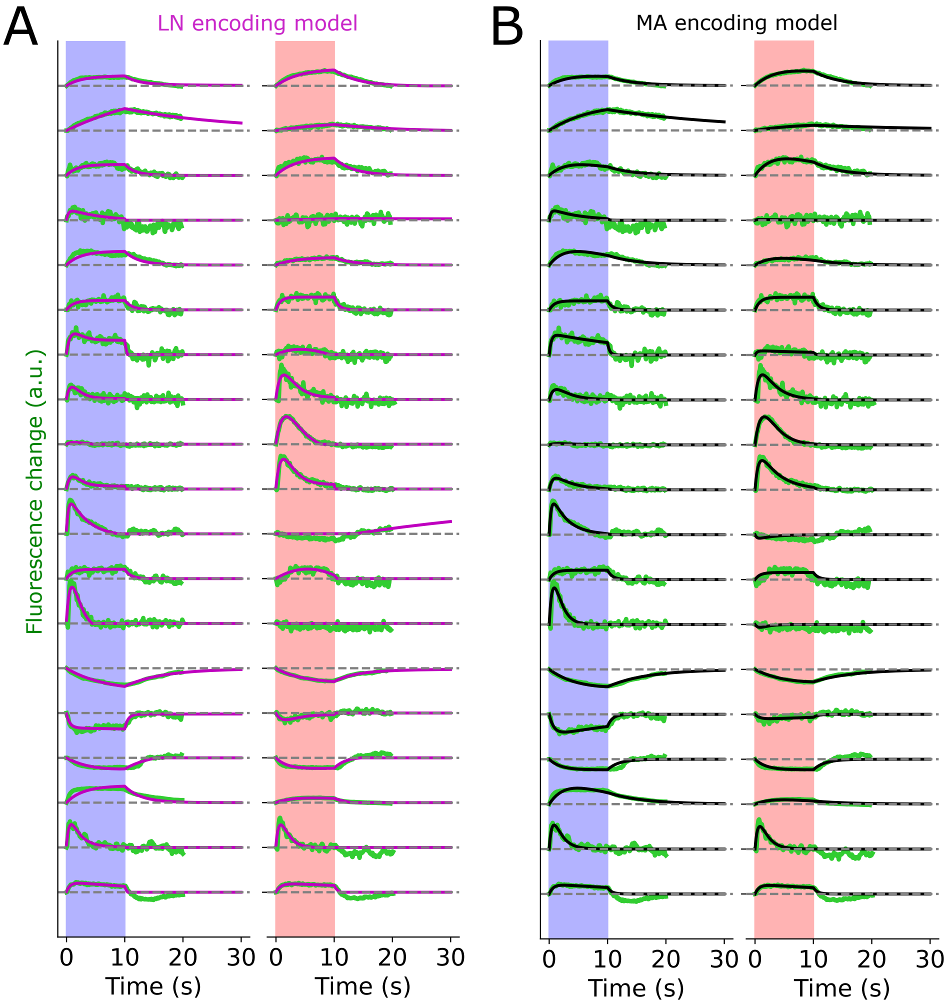


**Fig S2: Larger example set of encoding model fits to neural recordings.** A. Example MA fits (black) to song-element responses (green) recorded in Baker et al 2022 (1). B. As in A but for LN encoding models with filters parameterized by the MA parameters. Formatting as in Fig 1E.

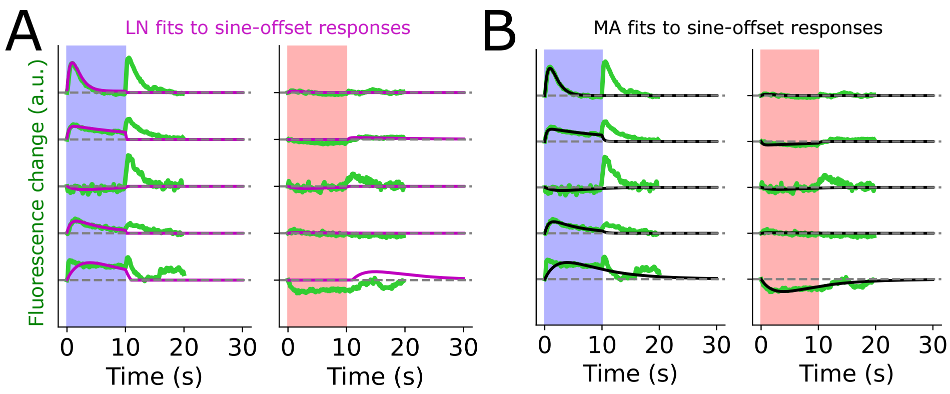


**Fig S3: Example sine-offset responses and model fits.** A. Example calcium responses and LN fits (magenta). B. As in A but with MA fits.

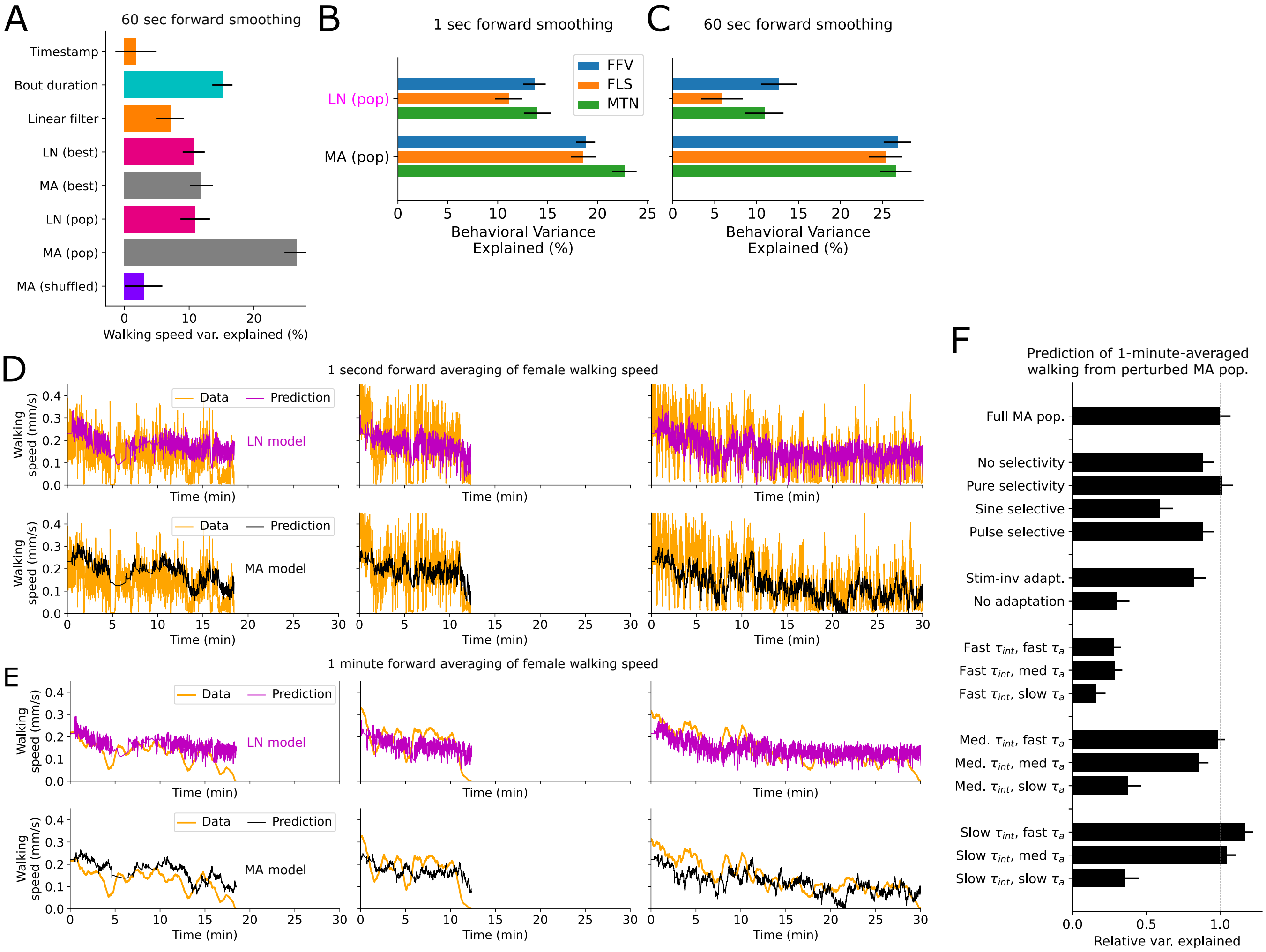


**Fig S4: Variance explained for different female behavioral variables and over different smoothing windows.** A. As in Fig 1J, but predicting female walking speed forward-averaged over a 60 s window. B. Comparison of female forward velocity (FFV), absolute female lateral speed (FLS) (relative to the female's body axis), and total female motion (MTN) i.e. walking speed, predictions using LN or MA populations, after forward-smoothing female behavior in a 1-second window. C. As in B but for a 60s window. D. Additional examples of walking speed predictions from the LN vs MA population models for three different held-out trials (as in Fig 1H). E. As in D, but using a 1-minute forward averaging window for female walking speed. F. As in Fig 2A, but for 1-minute-forward-averaged female walking speed.

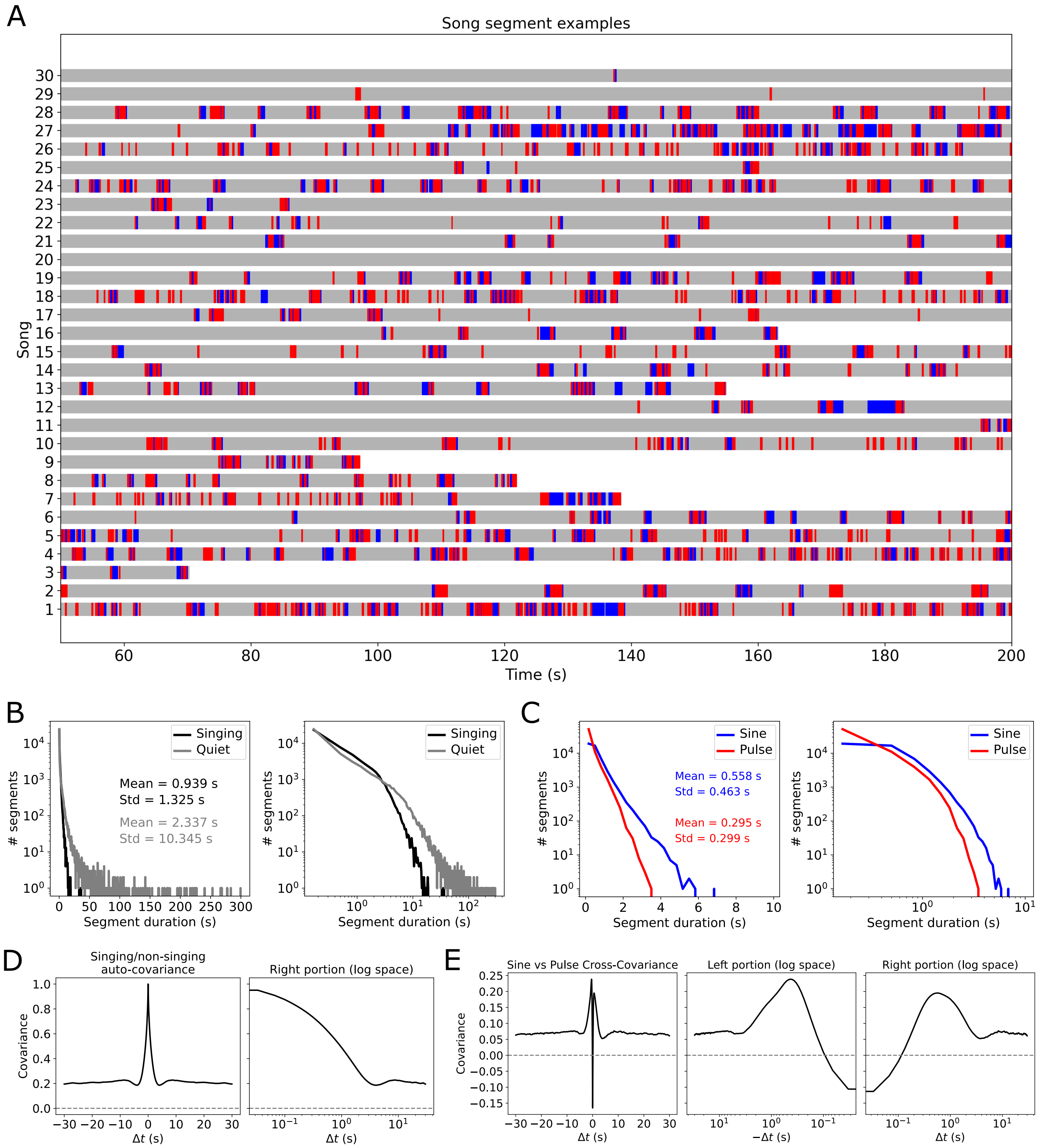


**Fig S5: Song examples and statistics.** A. Example songs; each row corresponds to a courtship ritual with a unique pair of virgin flies. Gray = quiet, blue = sine song, red = pulse song. Songs ending before 200s indicate that copulation occurred. B. Duration distribution of contiguous singing or quiet segments. Left plot is semi-log, right plot is log-log. C. As in D but for contiguous sine or pulse segments. D. Autocovariance function of singing vs non-singing, computed by aggregating all songs without normalization. The apparent nonzero offset likely partly reflects individual variability. E. Cross-covariance functions between binarized sine and pulse song, computed by aggregating all songs.

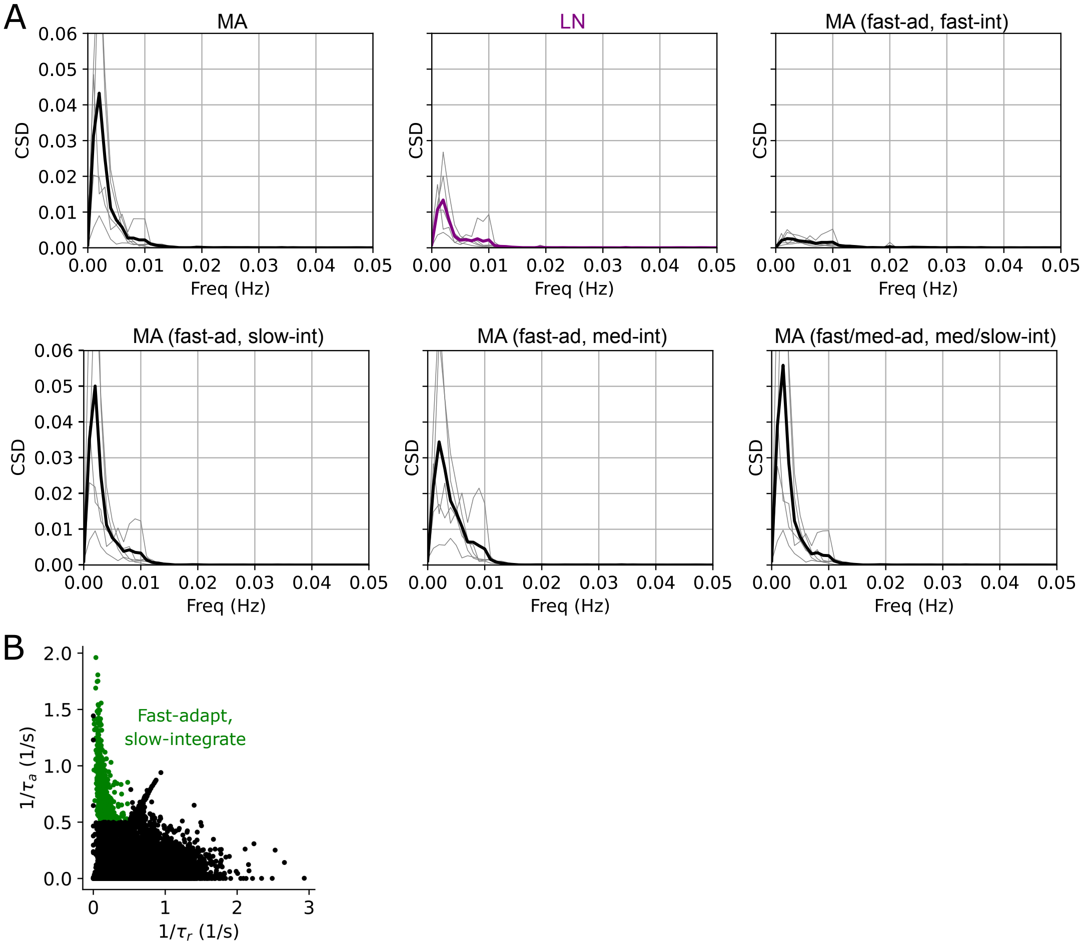


**Fig S6: Target/prediction alignment for different encoding models and fast-adapt slow-integrate region.** A. Cross-spectral density between target (female walking speed, 1-s forward smoothing) and prediction for held-out courtship sessions, for 5 training/test set splits (thick line shows average). The final panel (fast/med-ad, med/slow-int) uses $\tau_a$ sampled uniformly from 100 ms to 20 s and $\tau_{int}$ from 2 to 120 s. B. MA fit parameters using Pacheco et al (2021) pan-neuronal imaging data (2), with fast-adapt-slow-integrate regime highlighted.

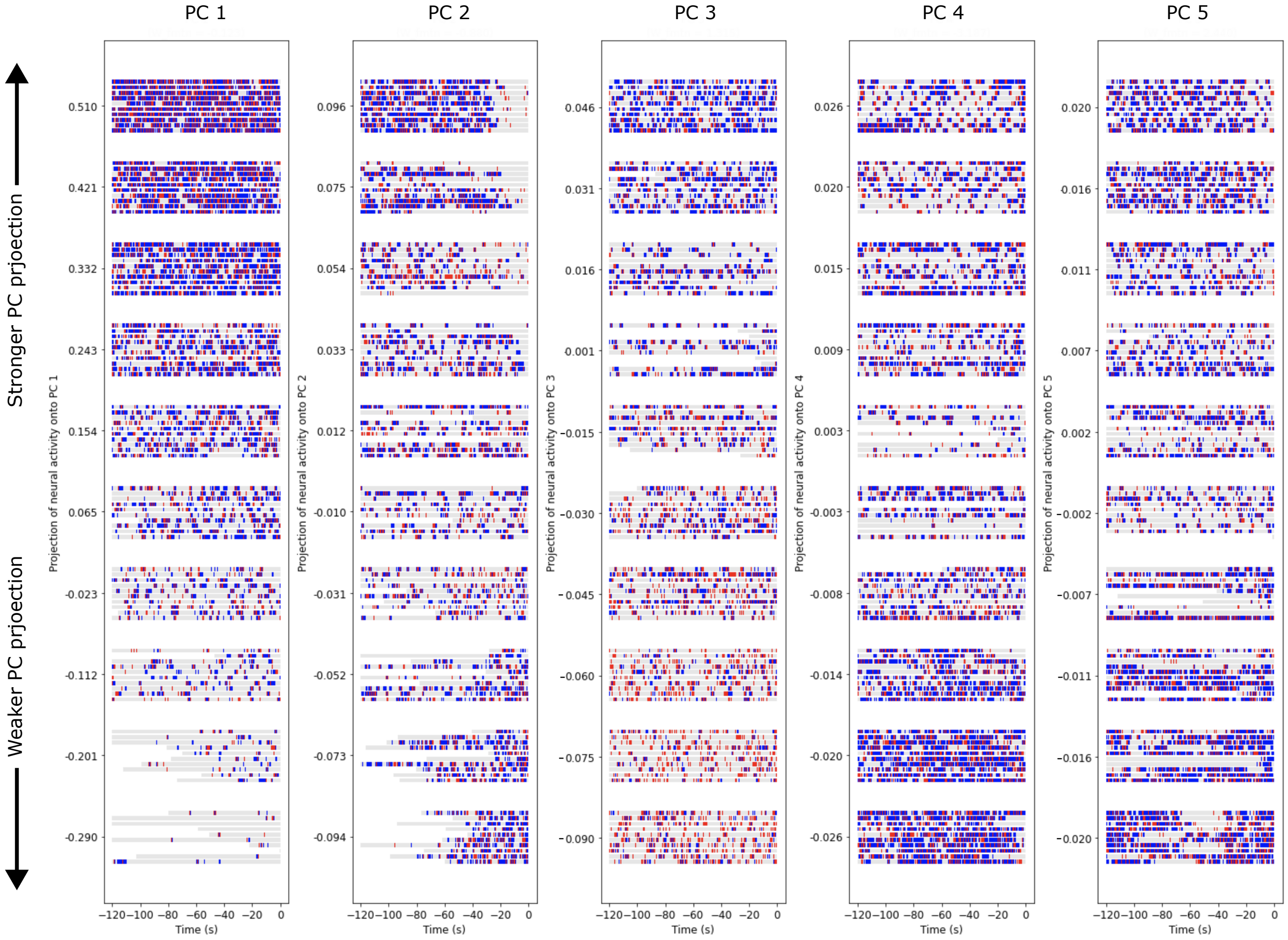


**Fig S7: Example songs driving activity along top 5 neural PCs of the 20-neuron fast-adapt-slow-integrate MA code.** Values of model neural activity projected onto each PC was split into deciles (using all timepoints across all songs). For each decile of each PC 10 songs/timepoints across the dataset were randomly sampled, and the last 2 minutes of song preceding that timepoint were plotted.

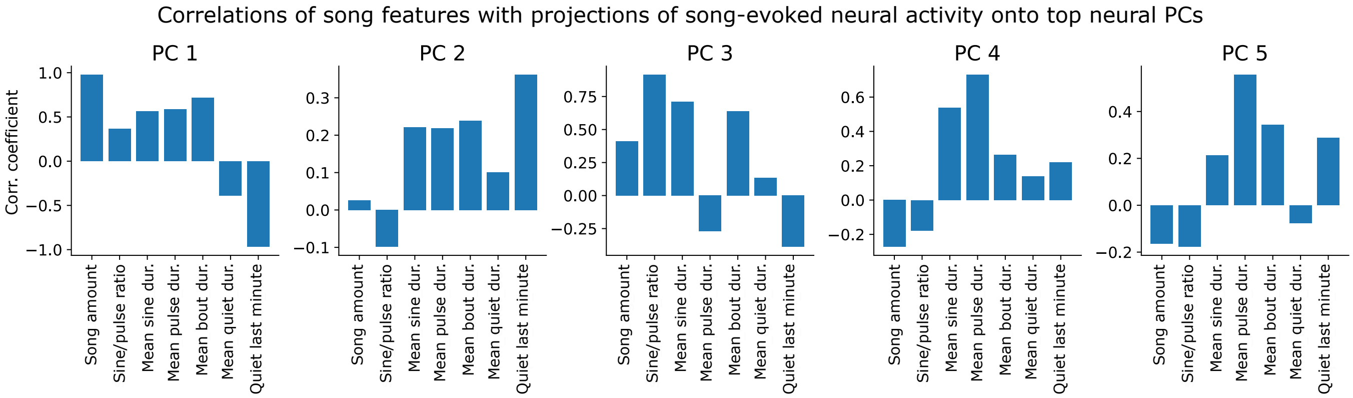


**Fig S8: Neural PCs interpreted in terms of song features**. For each song the values of several hand-picked features were computed. These were then correlated with projections of song-evoked MA population activity onto the top 5 neural PCs.

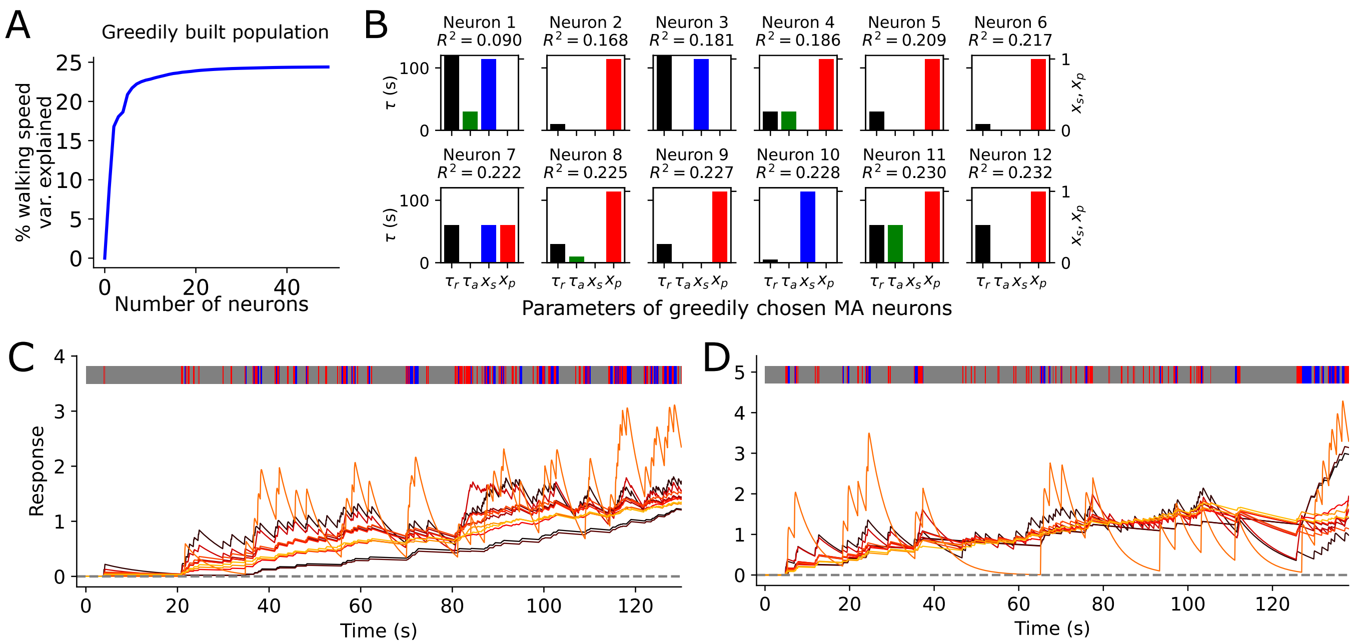


**Fig S9: Greedily constructed MA population.** A. Female walking speed variance (1-second forward averaging) explained in held-out courtship sessions from small population of MA neurons built up by greedily or iteratively choosing the next most predictive neuron to included in the population. B. Parameters of top 12 neurons recruited in the greedily constructed population in A. C. Population response to an example song segment. Each trace is the time-course of one neuron. D. As in C but for a different song.

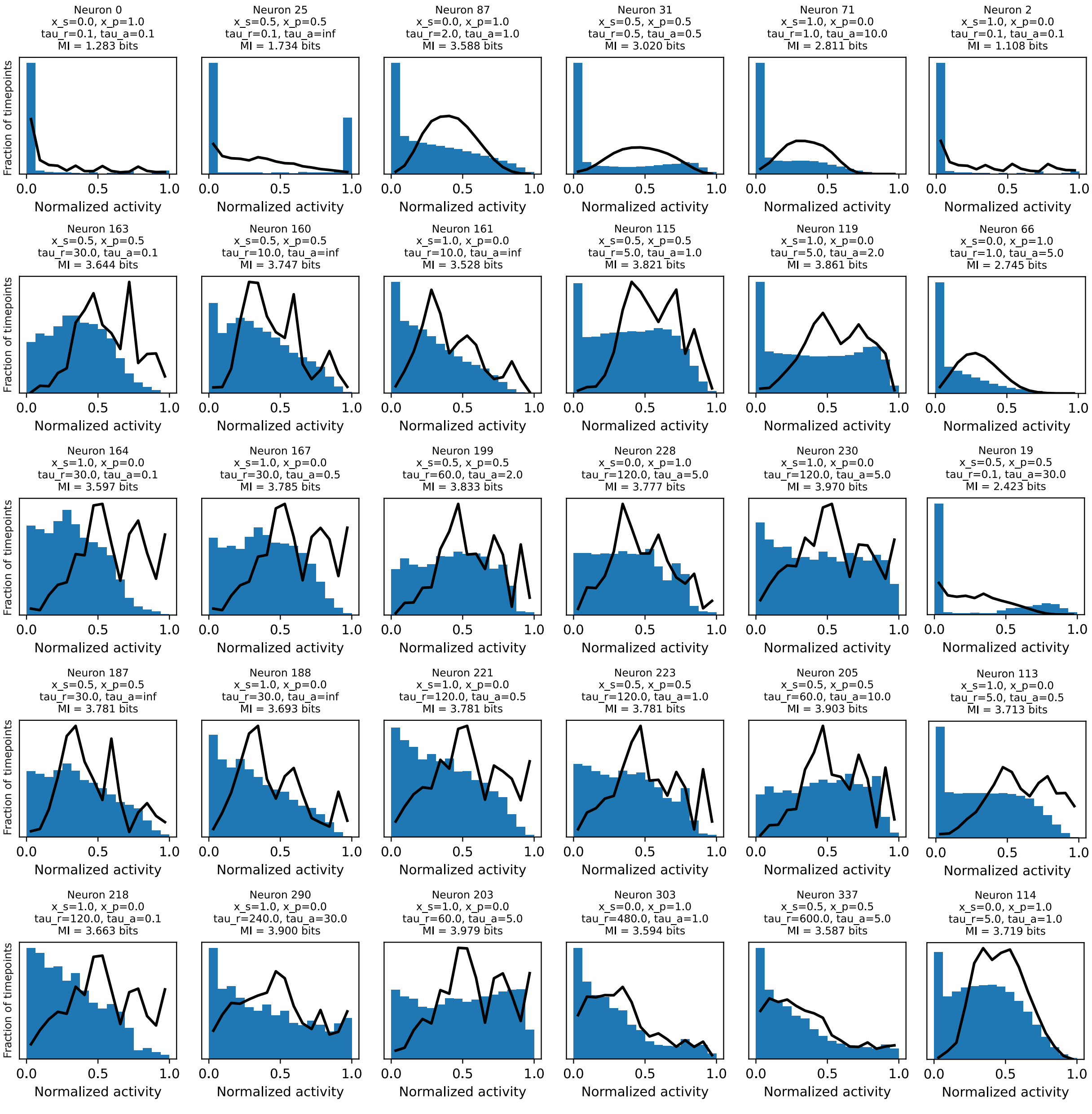


**Fig S10: Activity distributions of additional example model MA neurons.** Blue histograms show responses to natural song, binned into 16 bins. Black traces are histograms of responses to scrambled song. Activity is normalized to maximum observed response.

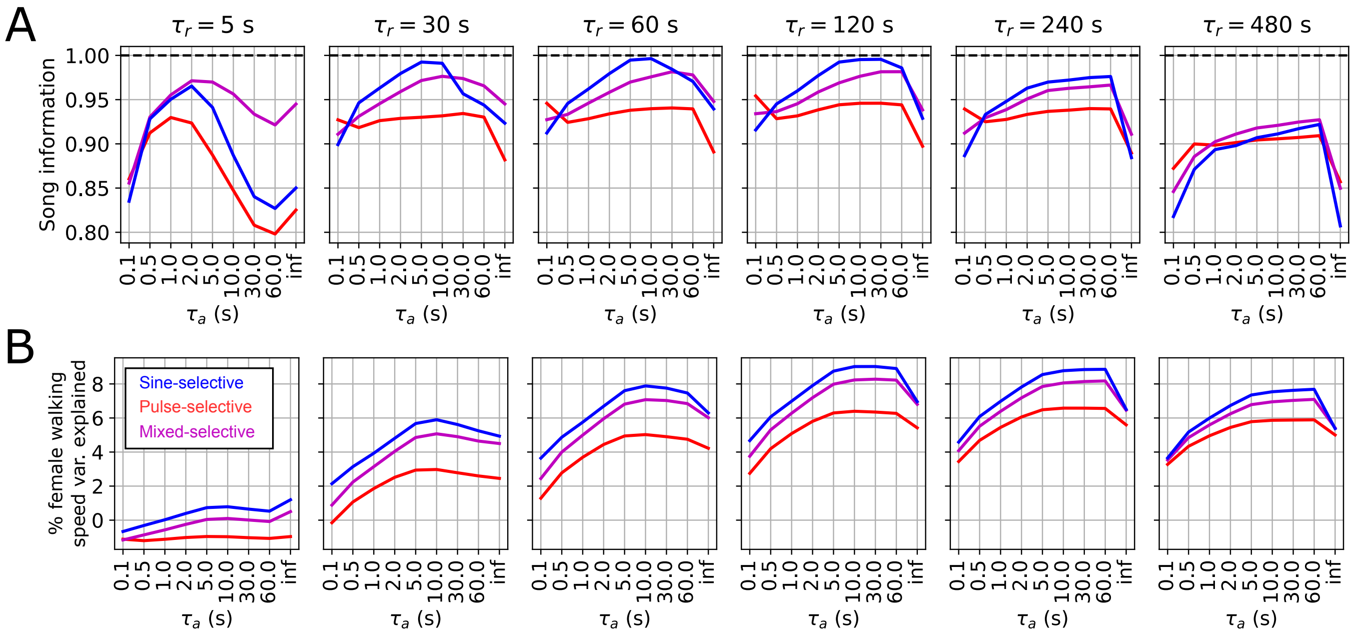


**Fig S11: Song information and walking speed variance explained by single MA neurons.** A. Mutual information between song and single MA neuron responses as a function of MA neuron parameters. Each panel shows a set of neurons with the indicated $\tau_{int}$. Blue lines indicate sine-selective neurons ($x_s = 1, x_p = 0$), red lines pulse-selective neurons ($x_s = 0, x_p = 1$) and magenta lines mixed-selective neurons ($x_s = x_p = 0.5$). B. As in A except depicting female walking speed variance explained by any single MA neuron, vs the MA neuron parameters.

 Rich Pang, Christa Baker, Mala Murthy, and Jonathan Pillow

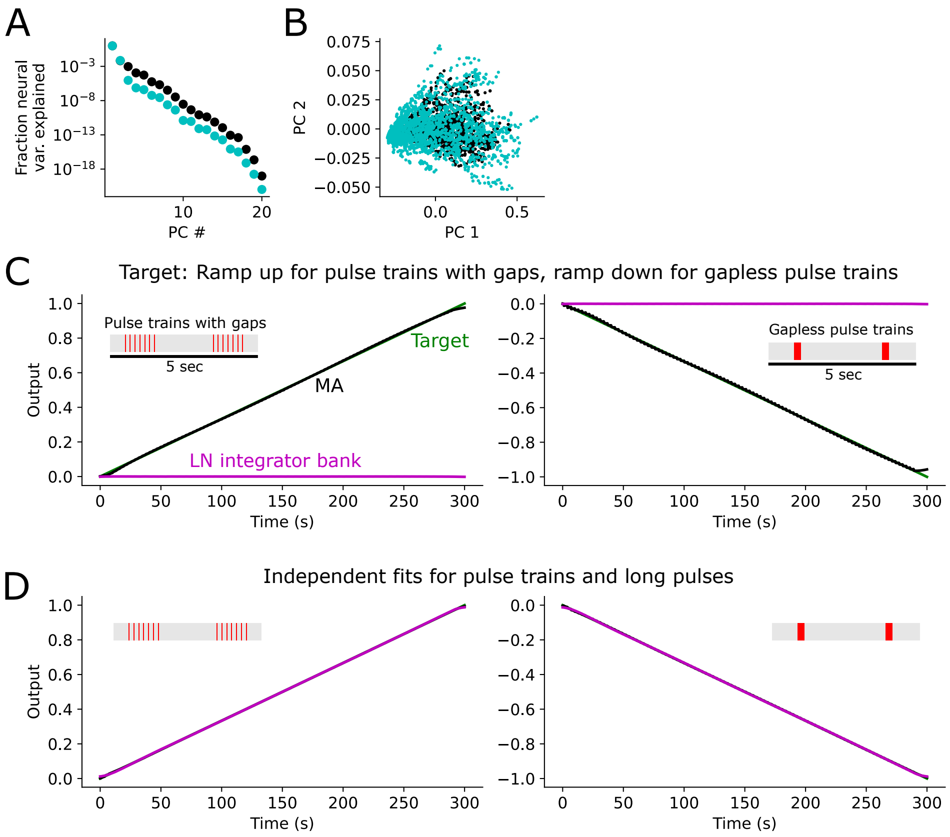


**Fig S12: Linear projections of song-evoked model neural population responses.** A. Fraction of variance explained by principal components of population response to example natural song (black) segments vs song segments that have been scrambled to remove temporal correlations (cyan). B. Song-evoked responses projected onto top 2 PCs for natural vs scrambled songs. C. Linear projections of MA vs LN integrator bank trained (Ridge Regression) to reproduce an increasing response to trains of 5 pulses interleaved with gaps (left) and a decreasing response to trains of 5 contiguous pulses. Here the MA model almost perfectly matches the target, unlike the LN model. This means that the MA model can distinguish pulse trains with vs without gaps, but the LN model cannot (because of its slow time constant). D. As in C except with independent training to each scenario. Here both the MA and LN model match the target.

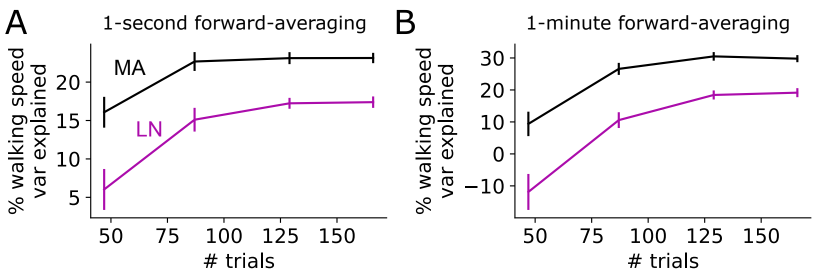


**Fig S13: Female walking speed variance explained vs number of trials.** A. Walking speed variance explained using artificial neural recordings generated by MA and LN population encoding models (LN model fit using the MA-step-response-matching procedure), using 1-second forward averaging of female walking speed. Error bars show standard error over 30 random train/test splits. B. As in A but using 1-minute forward averaging of female walking speed. In each plot the first 47 trials came from the male NM91 strain, the next 40 from ZH23, the next 42 from CarM03, and the final 37 from ZW109 (3).

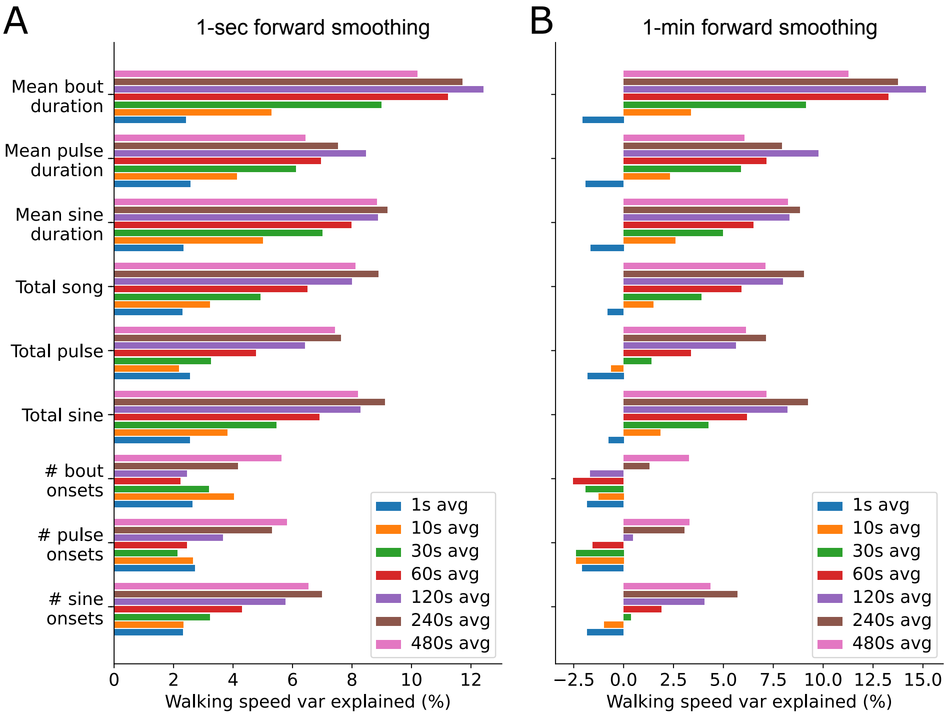


**Fig S14: Prediction of female locomotion from hand-picked song features**. A. Female walking speed variance explained (using 1-second forward smoothing) from a range of song features, estimated over several windows preceding the timepoint of the prediction. Variance explained is computed by averaging over 30 training/test splits of the data. B. As in A but using 1-minute forward smoothing.

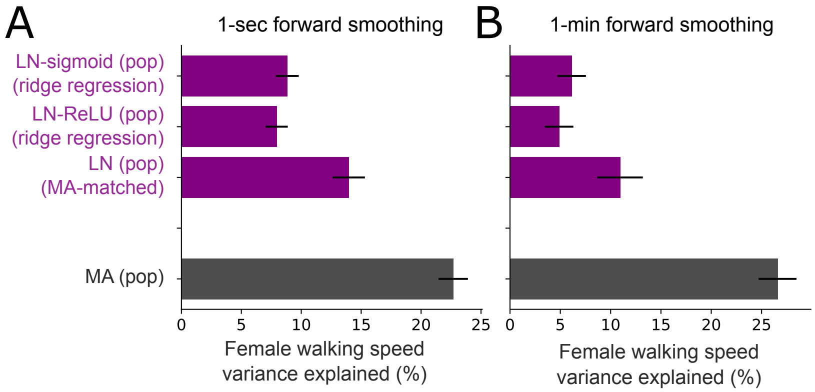


**Fig S15: Prediction of female locomotion from LN encoding population fit to neural data using ridge regression.** A. Female walking speed variance explained by different versions of the LN model, and from the MA model. LN-sigmoid: LN model fit with ridge regression using sigmoid nonlinearity. LN-ReLU: LN model fit with ridge regression using ReLU nonlinearity. B. As in A but predicting female walking speed forward-smoothed with a 1-minute window. This figure shows that the MA-matching procedure for fitting the LN model yielded more conservative results (better female walking speed prediction) than using ridge regression.

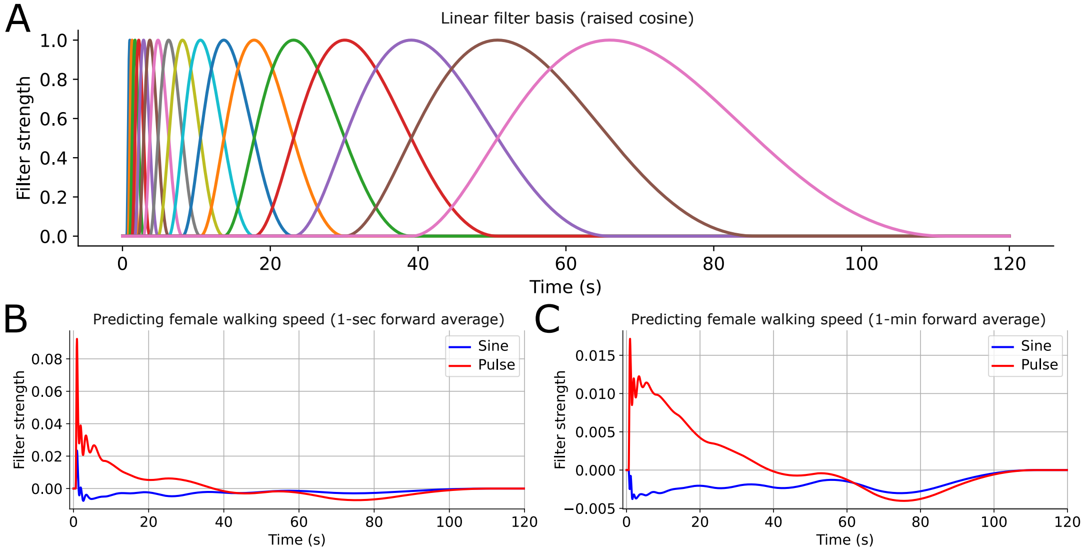


**Fig S16: Song-to-female-locomotion linear filter.** A. Basis functions (17 raised cosine functions). B. Sine- and pulse-filter reconstruction predicting female walking speed processed via a 1-second forward averaging window. C. As in B but for using a 1-minute forward averaging window.

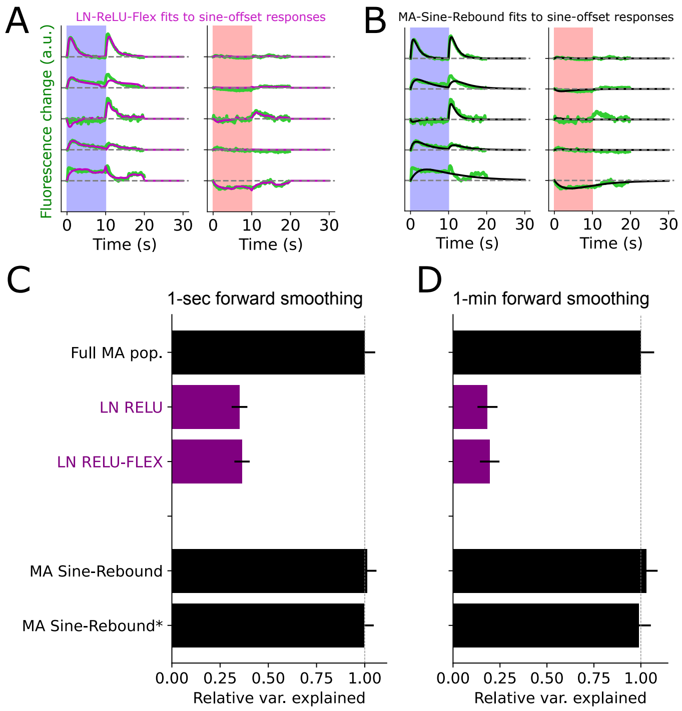


**Fig S17: Example sine-offset responses and model fits by MA-Sine-Rebound and LN-ReLU-Flex models.** A. Example sine-offset responses and fits with the "'LN-Relu-Flex" model, in which the nonlinearity was a piecewise linear function that was not constrained to be strictly monotonic. B. As in A but for an MA model in which quiet was treated as its own song mode. C. As in Fig 1J and 2A, predicting female walking speed smoothed over 1-second into the future. D. As in A but using a 1-min smoothing window. MA-Sine-Rebound* refers to model responses that treat pulse segments that follow sine as an additional unique song mode.
